# *U2AF1* is a haplo-essential gene required for cancer cell survival

**DOI:** 10.1101/2020.06.20.151035

**Authors:** Brian A. Wadugu, Amanda Heard, Sridhar N. Srivatsan, Michael O. Alberti, Matthew Ndonwi, Sarah Grieb, Joseph Bradley, Jin Shao, Tanzir Ahmed, Cara L. Shirai, Ajay Khanna, Dennis L. Fei, Christopher A. Miller, Timothy A. Graubert, Matthew J. Walter

**Author notes:** **Corresponding author**: Matthew Walter, Campus Box 8007, Washington University School of Medicine, 660 South Euclid Avenue, St. Louis, MO 63110.

## Abstract

Somatic mutations in the spliceosome gene *U2AF1* are common in patients with myelodysplastic syndromes. *U2AF1* mutations that code for the most common amino acid substitutions are always heterozygous, and the retained wild-type allele is expressed, suggesting that mutant hematopoietic cells may require the residual wild-type allele to be viable and cause disease. We show that hematopoiesis and RNA splicing in *U2af1* heterozygous knock-out mice was similar to control mice, but that deletion of the wild-type allele in U2AF1(S34F) heterozygous mutant expressing hematopoietic cells (i.e., hemizygous mutant) was lethal. These results confirm that U2AF1 mutant hematopoietic cells are dependent on the expression of wild-type U2AF1 for survival *in vivo* and that *U2AF1* is a haplo-essential cancer gene. Mutant U2AF1 (S34F) expressing cells were also more sensitive to reduced, but not absent, expression of wild-type U2AF1 than non-mutant cells. Furthermore, mice transplanted with leukemia cells expressing mutant U2AF1 had significantly reduced tumor burden and improved survival after the wild-type *U2af1* allele was deleted compared to when it was not deleted. These results suggest that selectively targeting the wild-type *U2AF1* allele in heterozygous mutant cells could induce cancer cell death and be a therapeutic strategy for patients harboring *U2AF1* mutations.

## Introduction

Myelodysplastic syndromes (MDS), the most common myeloid cancers in adults in the United States, are characterized by low peripheral blood counts in patients and a propensity to progress to secondary acute myeloid leukemia (AML) (1, 2). Somatic mutations in spliceosome genes (e.g., *SF3B1, SRSF2, U2AF1,* and *ZRSR2*) occur in up to 50% of MDS patients, making them the most common group of genes mutated in MDS (3–9). U2AF1 is a U2 auxiliary factor protein that recognizes and binds the AG dinucleotide in the intronic 3’ splice acceptor site of pre-mRNA (10). *U2AF1* mutations impact the RNA binding zinc finger domains in up to 11% of MDS patients, with the most frequent coding for the U2AF1(S34F) substitution (9, 11). *U2AF1* mutations are always heterozygous with the residual wild-type allele expressed, suggesting that mutant hematopoietic cells may require one copy of wild-type *U2AF1* for survival. Consistent with this, *U2AF1* has been nominated as a haplo-essential cancer gene (e.g., cancer cells with a *U2AF1* heterozygous mutation are under selective pressure to maintain at least one copy of the *U2AF1* wild-type allele) based on analysis of large-scale sequencing studies (12).

New treatment strategies are emerging for MDS that exploit vulnerabilities present in spliceosome mutant cells but not in normal cells. As an example, our group and others reported that spliceosome mutant hematopoietic cells are more sensitive to drugs that modulate splicing compared to wild-type cells (13–16). Because haplo-essential cancer genes are predicted to be dependent on the expression of the residual wild-type allele for cell survival, this could provide another vulnerability to target in U2AF1 mutant cells, as has been observed for mutant SRSF2 cells in mice (14). *In vitro* studies have shown that expression of the wild-type allele was required for survival of an immortalized human bronchial epithelial cell line harboring U2AF1(S34F) at the endogenous locus (17). Knowing if *U2AF1* is a haplo-essential gene required for hematopoietic cell survival *in vivo* could provide insight into MDS disease pathogenesis and have treatment implications for cancers harboring a heterozygous mutation.

In this study, we used *U2af1* heterozygous and homozygous knock-out mice to define the role of wild-type U2AF1 in hematopoiesis and determine whether mutant U2AF1-expressing hematopoietic cells, including leukemia cells, require the expression of the residual wild-type *U2af1* allele for cell survival. We find that modulating the expression ratio of wild-type to mutant U2AF1 may be a therapeutic approach to selectively kill U2AF1 mutant expressing cancer cells *in vivo*.

## Results

### Generation of a conditional *U2af1* knock-out mouse

We generated a *U2af1* knock-out (KO) mouse model to study the role of U2AF1 in hematopoiesis and test whether *U2AF1* is a haplo-essential cancer gene (i.e., requiring the expression of at least one copy of the wild-type allele for cell viability). Constitutive deletion of U2AF1 is lethal in yeast, *C. elegans*, drosophila, and zebrafish (18–21). Therefore, we created a conditional *U2af1* knock-out (*U2af1* KO) allele in mice to study the hematopoietic-specific effects of *U2af1* deletion. To create the conditional *U2af1* KO allele, we flanked exon 2 with loxP sites in wild-type C57B/L6 embryonic stem cells (ES cells) (**Figure 1A, Supplemental Figure 1A**). Exon 2 is common in all *U2af1* isoforms, codes for one of the two zinc finger domains critical for binding RNA, and contains the sequence that codes for the S34F substitution (11). Deletion of exon 2 is ideal for creating the *U2af1* KO mouse because it results in a frameshift and predicted early termination codon when exon 1 splices into downstream exons (except when exon 1 splices into exon 7 or 8 and potentially results in a protein lacking both zinc finger domains and the U2AF homology motif). We confirmed correct targeting of the *U2af1* locus in five embryonic stem (ES) cell clones by Southern blotting and PCR (**Figure 1B, Supplemental Figure 1B, C**). Whole-genome sequencing further validated the correct targeting of the locus (**Supplemental Figure 1D**). Mice harboring a *U2af1* floxed allele were crossed to existing C57BL/6 Cre-expressing lines, including Cre-*ERT2*, and the hematopoietic cell-specific Cre-expressing lines, *Mx1-*Cre and *Vav1-*Cre (22–24).

**Figure 1.**
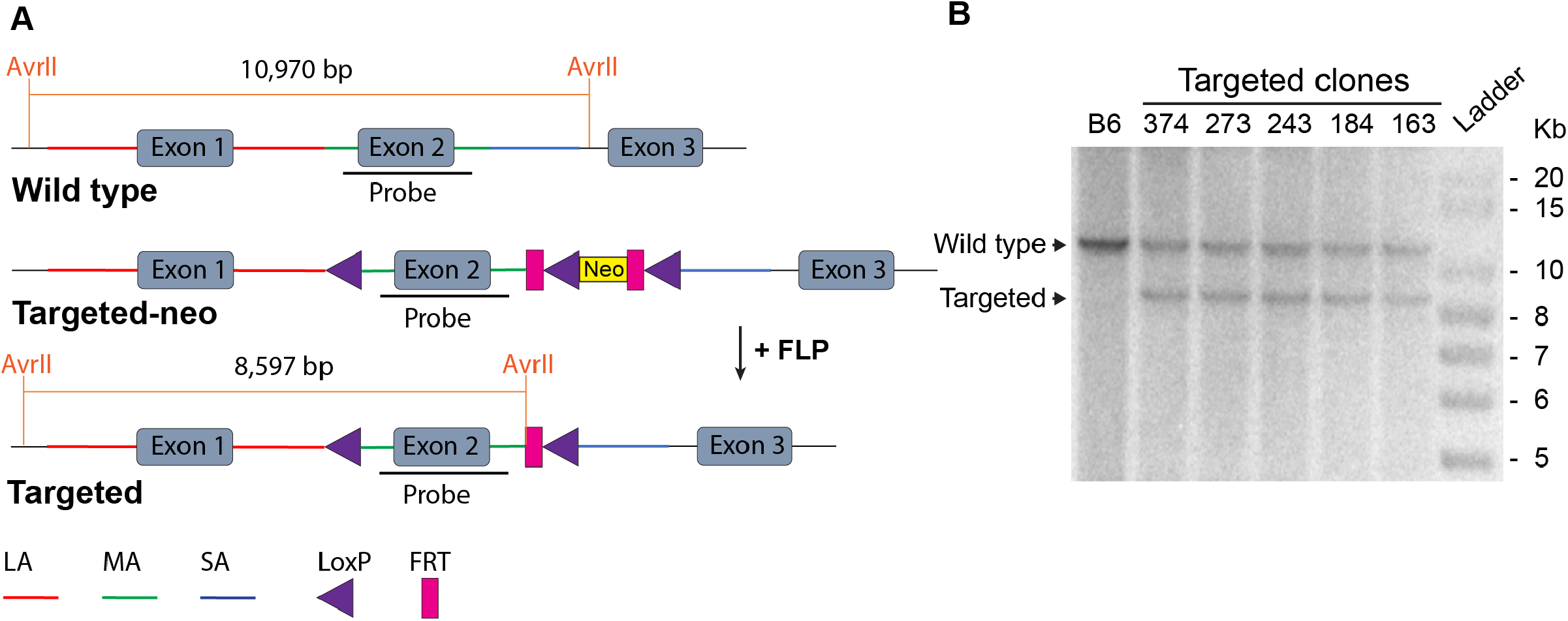
Generation of a conditional *U2af1* knock-out allele. (A) Targeting strategy to insert loxP sites flanking *U2af1* exon 2. (B) Successful targeting of five embryonic stem cell clones was verified by Southern blotting after digestion with the restriction enzyme AvrII. Wild-type C57BL/6 (B6) DNA was used as a control, and clone 243 was used to generate the *U2af1* knock-out mouse. LA (long arm), MA (middle arm), SA (Short arm).

### Embryonic expression of U2AF1 in hematopoietic cells is essential for normal hematopoiesis and viability

In order to understand the role of wild-type U2AF1 in normal embryonic hematopoiesis, we intercrossed *U2af1*^wt/flox^ mice with *Vav1*-Cre/*U2af1*^wt/flox^ mice and analyzed the frequency of observed versus expected genotypes at birth. Homozygous deletion of *U2af1* (*Vav1*-Cre/*U2af1*^flox/flox^) was embryonic lethal, evidenced by 100% lethality at birth, while the control *Vav1*-Cre and *U2af1*^wt/flox^ mice, as well as the heterozygous *U2af1* KO mice (*Vav1*-Cre/*U2af1*^wt/flox^), were born at expected frequencies (**Figure 2A, Supplemental Figure 2A**). These results indicate that hematopoietic expression of U2AF1 is essential for viability during embryonic development.

**Figure 2.**
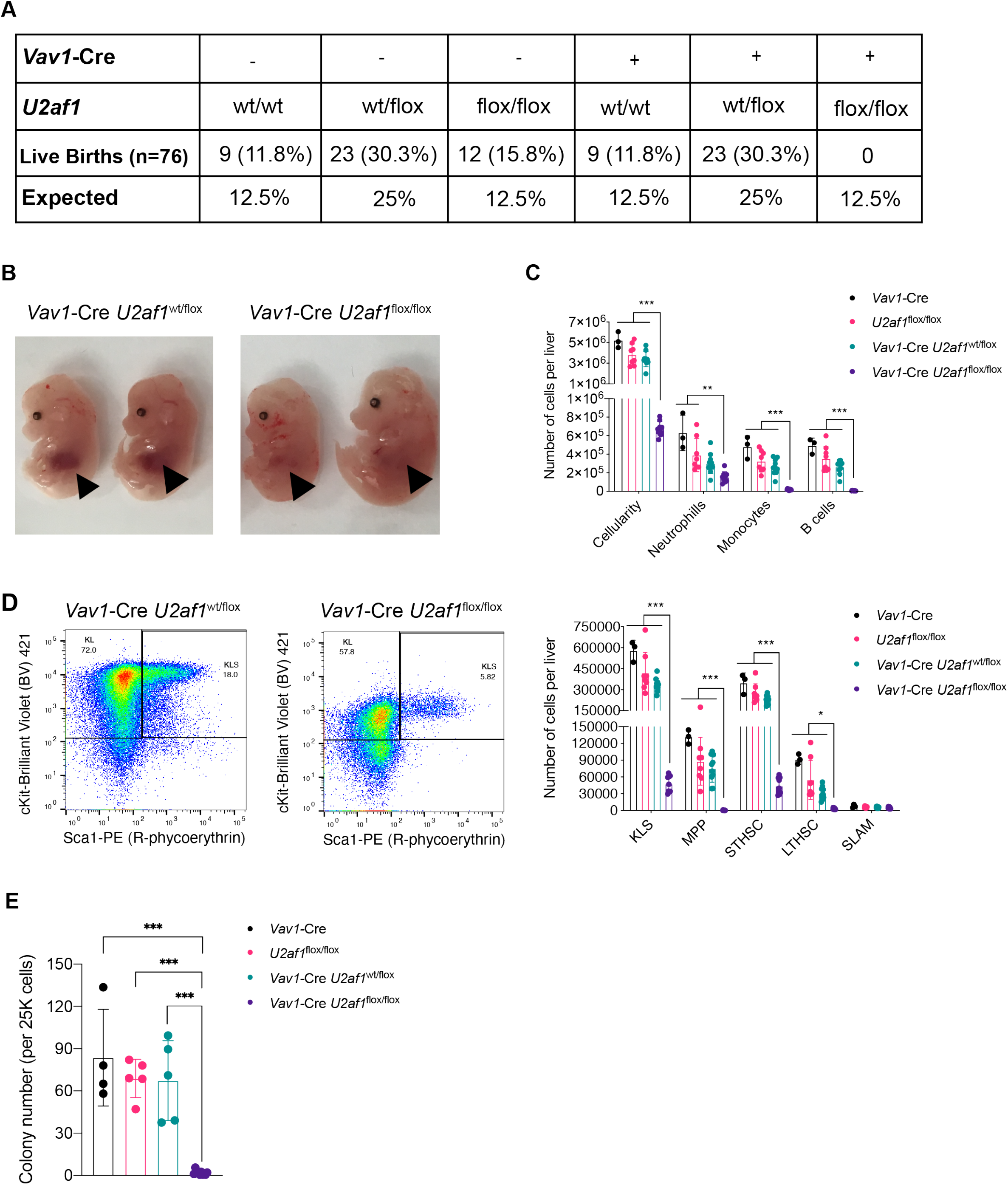
Embryonic deletion of *U2af1* in hematopoietic cells is lethal and reduces the number of myeloid, lymphoid, and hematopoietic stem and progenitor cells. (A) *U2af1*^wt/flox^ mice were intercrossed with *Vav1*-Cre/*U2af1*^wt/flox^ mice, and the frequency of observed versus expected genotypes were analyzed at birth (Chi squared=11.895, 5 degrees of freedom, two-tailed p=0.0363, n=76). (B) Comparison of fetal liver development of embryos at embryonic day 14.5 (E14.5) between *Vav1*-Cre/*U2af1*^wt/flox^ and *Vav1*-Cre/*U2af1*^flox/flox^ mice. (C) E14.5 fetal liver hematopoietic cellularity, and absolute counts of neutrophils, monocytes, and B cells in the fetal liver of the *Vav1*-Cre, *U2af1*^wt/flox^, and *Vav1*-Cre/*U2af1*^wt/flox^ embryos were compared to *Vav1*-Cre/*U2af1*^flox/flox^ embryos (n=3-10). (D) The proportion of progenitor cells (KL and KLS) of *Vav1*-Cre/*U2af1*^wt/flox^ compared to *Vav1*-Cre/*U2af1*^flox/flox^ mice (representative flow figures on the left), and the absolute numbers of hematopoietic stem and progenitor cells (KLS, MPP, STHSC, LTHSC, SLAM) in the fetal liver of *Vav1*-Cre, *U2af1*^wt/flox^, and *Vav1*-Cre/*U2af1*^wt/flox^ embryos compared to *Vav1*-Cre/*U2af1*^flox/flox^ embryos (n=3-10). (E) Colony numbers from 25,000 bulk fetal liver cells isolated from *Vav1*-Cre, *U2af1*^wt/flox^, and *Vav1*-Cre/*U2af1*^wt/flox^ embryos compared to *Vav1*-Cre/*U2af1*^flox/flox^ embryos after 8 days (n=4-7). KL (Lineage^−^/cKit^+^/Sca1^−^), KLS (Lineage^−^/cKit^+^/Sca1^+^). KLS subpopulations: SLAM (KLS/CD150^+^/CD48^−^), multipotent progenitors (MPP; KLS/CD34^+^/Flk2^+^), short-term stem cells (ST-HSC; KLS/CD34^+^/Flk2^−^), long-term stem cells (LT-HSC; KLS/CD34^−^/Flk2^−^). All data are presented as mean +/− SD. *p<0.05, **p<0.01, ***p<0.001 by one-way ANOVA with Tukey’s multiple comparison test.

Because embryonic-induced homozygous deletion of *U2af1* using *Vav1*-Cre was lethal prior to birth, we studied hematopoiesis at embryonic day 14.5 (E14.5) when the expected frequency of genotypes was observed (data not shown). Comparison of fetal liver development of embryos at E14.5 showed that the homozygous *U2af1* KO mice had underdeveloped livers compared to the heterozygous *U2af1* KO mice (**Figure 2B**) and the two control genotypes (**Supplemental Figure 2B**). Analysis of the fetal liver hematopoietic cells at E14.5 revealed that embryonic homozygous deletion of *U2af1* in *Vav1*-Cre/*U2af1*^flox/flox^ embryos led to a reduction in cellularity, neutrophil, monocyte and B cell numbers, while the heterozygous *U2af1* KO and the two control genotypes were similar and unaffected (**Figure 2C**). Furthermore, analysis of flow cytometry data revealed that E14.5 homozygous KO (*Vav1*-Cre/*U2af1*^flox/flox^) fetal livers had a reduced proportion of progenitor cells (KL [Lineage^−^/cKit^+^/Sca1^−^], and KLS [Lineage^−^/cKit^+^/Sca1^+^]) compared to heterozygous KO (*Vav1*-Cre/*U2af1*^wt/flox^) fetal livers (**Figure 2D**, representative flow figures on the left). The absolute number of hematopoietic stem and progenitor cells (KL, KLS, common myeloid progenitors [CMP; KL/CD34^+^/Fcγ^−^], megakaryocyte erythroid progenitors [MEP; KL/CD34^−^/Fcγ^−^], granulocyte-macrophage progenitors [GMP; KL/CD34^+^/Fcγ^+^], multipotent progenitors [MPP; KLS/CD34^+^/Flk2^+^], short-term stem cells [ST-HSC; KLS/CD34^+^/Flk2^−^], long-term stem cells [LT-HSC; KLS/CD34^−^/Flk2^−^], but not SLAM cells [KLS/CD150^+^/CD48^−^]) were significantly reduced in the homozygous *U2af1* KO fetal liver compared to the heterozygous *U2af1* KO and two control genotypes (**Figure 2D, Supplemental Figure 2C**). In a methylcellulose colony-forming assay, we observed that the hematopoietic progenitor cells from homozygous *U2af1* KO embryos have significantly impaired colony-forming ability compared to the heterozygous *U2af1* KO and two control embryos (**Figure 2E**). Collectively, these results suggest that U2AF1 expression is essential for hematopoiesis and hematopoietic cell viability during embryonic development.

### Homozygous deletion of *U2af1* in adult mice results in multi-lineage bone marrow failure

To study the hematopoietic cell-intrinsic effects of *U2af1* deletion in adult mice, we generated homozygous *Mx1*-Cre/*U2af1*^flox/flox^, heterozygous *Mx1*-Cre/*U2af1*^wt/flox^, control *Mx1*-Cre, and control *U2af1*^flox/flox^ mice. First, we harvested bulk bone marrow cells from these donor mice and performed a non-competitive bone marrow transplant into lethally irradiated congenic recipient mice to study hematopoietic cell-intrinsic effects (**Figure 3A**). Following polyinosinic-polycytidylic acid (pIpC)-induced *U2af1* deletion in *Mx1-*Cre expressing mice, mice harboring homozygous *U2af1* KO cells, but not other genotypes (including mice with heterozygous *U2af1* KO cells), became moribund (**Supplemental Figure 3B**). Mice harboring homozygous *U2af1* KO cells developed pancytopenia as early as 8-11 days post pIpC-induced *U2af1* deletion compared to all other genotypes (**Figure 3B**). Homozygous deletion of *U2af1* also led to rapid bone marrow failure and a reduction in the absolute number of bone marrow and spleen neutrophils, monocytes, and B cells, but not T cells, 8-11 days post-pIpC (**Figure 3C, D, Supplemental Figure 3C, D**). Hematopoietic progenitor cells (KL and KLS) were also significantly reduced at 8-11 days post-pIpC (**Figure 3E, Supplemental Figure 3E**). The bone marrow failure was verified in a Cre-*ERT2* model. Bone marrow cellularity was reduced in Cre-*ERT2*/*U2af1*^flox/flox^ compared to the Cre-*ERT2* control mice three days post final dose of tamoxifen (**Figure 3F)**. These results show that wild-type U2AF1 is required for normal hematopoiesis and cell viability in adult mice.

**Figure 3.**
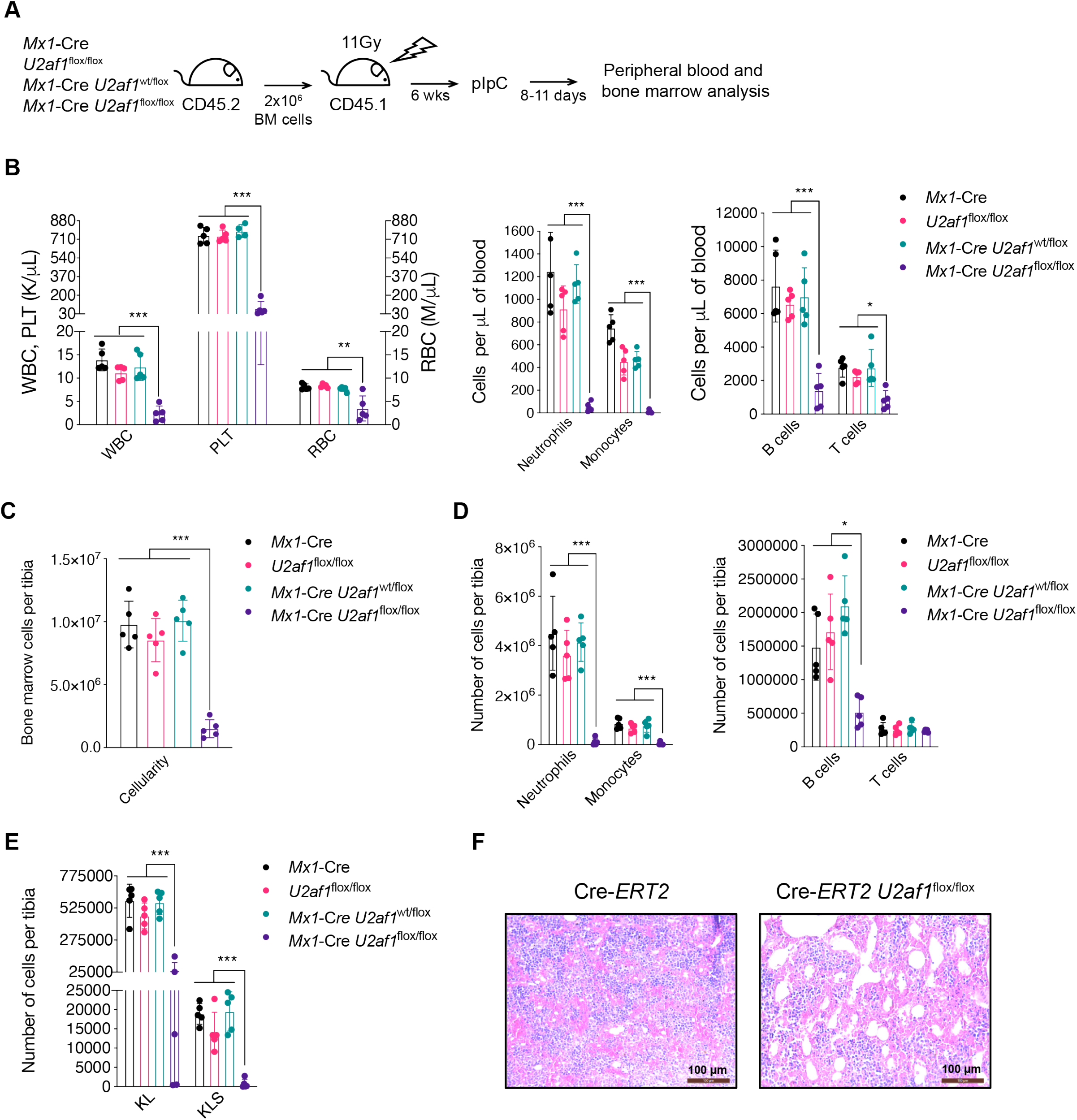
*U2af1* deletion induces multi-lineage bone marrow failure. (A) Experimental design of a non-competitive transplant of whole bone marrow cells from CD45.2 *Mx1*-Cre, *U2af1*^flox/flox^, *Mx1*-Cre*/U2af1*^wt/flox^, or *Mx1*-Cre/*U2af1*^flox/flox^ mice transplanted into lethally irradiated congenic wild-type CD45.1 recipient mice, followed by pIpC-induced Cre-activation and *U2af1* deletion. Analysis of the peripheral blood and bone marrow was done 8-11 days post pIpC. (B) Absolute numbers of peripheral white blood cells (WBC), platelets (PLT), red blood cells (RBC), neutrophils, monocytes, B cells and T cells of mice transplanted with whole bone marrow from *Mx1*-Cre, *U2af1*^wt/flox^, and *Mx1*-Cre/*U2af1*^wt/flox^ mice compared to *Mx1*-Cre/*U2af1*^flox/flox^ mice 8-11 days post pIpC. (C) Bone marrow cellularity of *Mx1*-Cre, *U2af1*^wt/flox^, and *Mx1*-Cre/*U2af1*^wt/flox^ mice compared to *Mx1*-Cre/*U2af1*^flox/flox^ mice 8-11 days post pIpC. (D) Total number of neutrophils, monocytes, B cells, and T cells per tibia at 8-11 days post pIpC for *Mx1*-Cre, *U2af1*^wt/flox^, and *Mx1*-Cre/*U2af1*^wt/flox^ mice compared to *Mx1*-Cre/*U2af1*^flox/flox^ mice. (E). Number of progenitor cells (KL [Lineage^−^/cKit^+^/Sca1^−^], and KLS [Lineage^−^/cKit^+^/Sca1^+^]) per tibia of pIpC-treated mice transplanted with *Mx1*-Cre, *U2af1*^wt/flox^, and *Mx1*-Cre/*U2af1*^wt/flox^ bone marrow cells compared to *Mx1*-Cre/*U2af1*^flox/flox^ bone marrow cells 8-11 days post pIpC. (F) H&E staining of bone marrow from Cre-ERT2/*U2af1*^flox/flox^ and Cre-ERT2 control mice at three days post-tamoxifen. The experimental design is similar to that in panel (A) with tamoxifen used for induction of *U2af1* deletion. All data are presented as mean +/− SD. *p<0.05, **p<0.01, ***p<0.001 by one-way ANOVA with Tukey’s multiple comparison test, n=5 per genotype.

### Hematopoietic stem cells are dependent on *U2af1* expression for survival and normal function

In order to study the hematopoietic cell-intrinsic effects of *U2af1* deletion on stem cell function, we performed a competitive repopulation transplant. We created mixed bone marrow chimeras using test hematopoietic cells from *Mx1*-Cre, *U2af1*^flox/flox^, *Mx1*-Cre/*U2af1*^wt/flox^ or *Mx1*-Cre/*U2af1*^flox/flox^ donor mice mixed with equal numbers of competitor cells from congenic wild-type donor mice (**Figure 4A**). As early as ten days following *Mx1*-Cre-induction by pIpC to delete *U2af1*, we observed a significant decrease in peripheral blood white blood cell chimerism (**Figure 4B**) in homozygous *U2af1* KO cells but not heterozygous *U2af1* KO or control cells in congenic recipient mice; this included complete loss of neutrophils and monocytes (**Figure 4C, D**), and a severe reduction in B cells and T cells (**Figure 4E, F**). Flow cytometric analysis of bone marrow and spleen chimerism ten months post-deletion of *U2af1* revealed a complete loss of homozygous *U2af1* KO bone marrow hematopoietic neutrophils and monocytes, as well as a severe reduction in B cells and T cells (**Figure 4G, Supplemental Figure 4B**). Furthermore, there was a complete loss of *U2af1* homozygous KO stem and progenitor cells (KL, KLS, CMP, GMP, MEP, MPP, ST-HSC, LT-HSC, and SLAM) in the mice bone marrow (**Figure 4H, Supplemental Figure 4C**). Collectively, these data indicate that multi-lineage hematopoiesis, stem cell function, and hematopoietic cell survival are dependent on U2AF1 expression, and that *U2af1* heterozygous KO cells that retain one *U2af1* allele are normal, with albeit a very mild disadvantage in T cell chimerism, at 120 days post-pIpC (**Figure 4F**).

**Figure 4.**
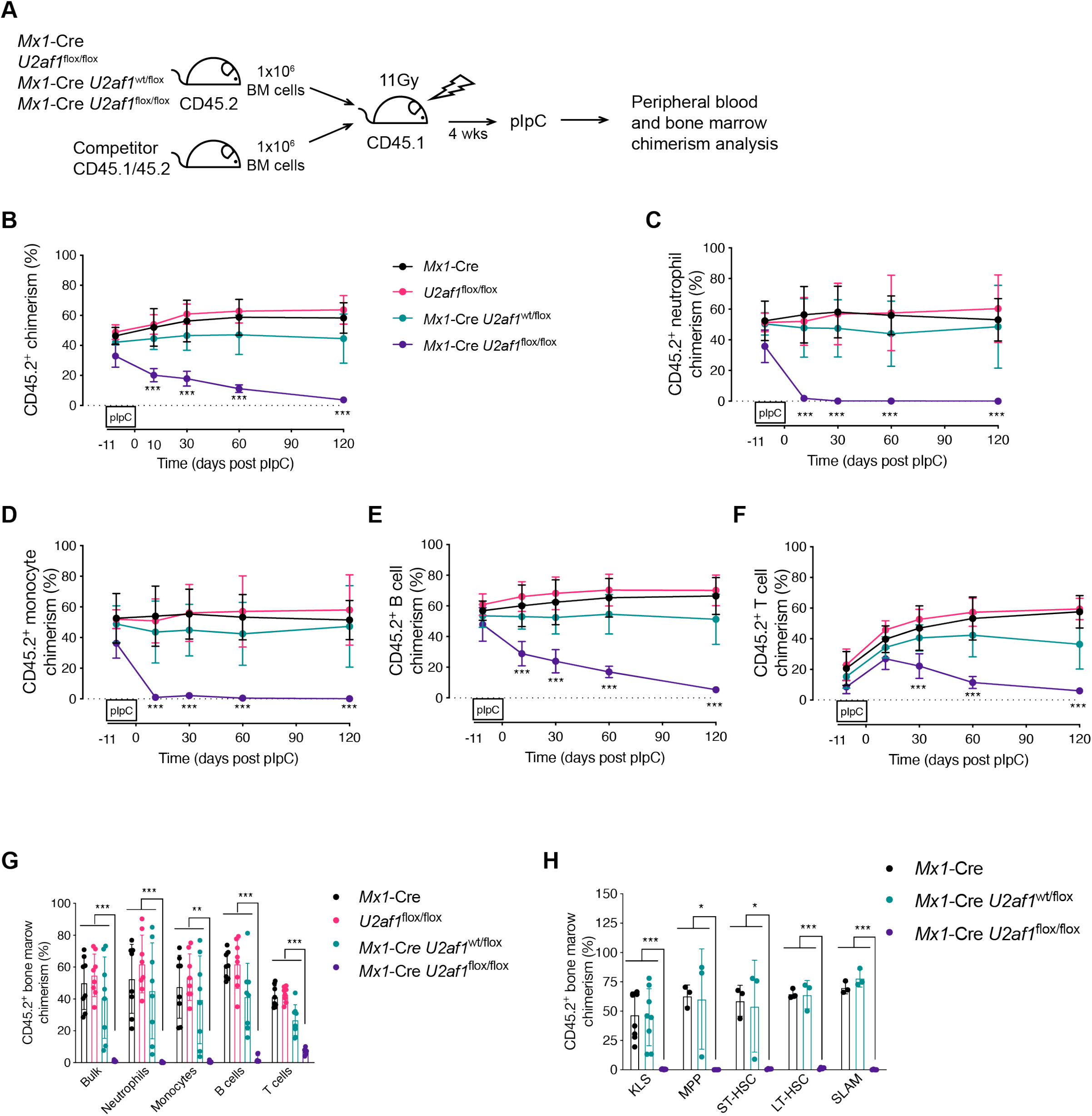
Hematopoietic stem cells are dependent on *U2af1* expression for survival. (A) Experimental design of a competitive transplant of whole bone marrow cells from CD45.2 *Mx1*-Cre, *U2af1*^flox/flox^, *Mx1*-Cre/*U2af1*^wt/flox^, or *Mx1*-Cre/*U2af1*^flox/flox^ mice mixed at a 1:1 ratio with congenic wild-type CD45.1/45.2 competitor cells followed by transplantation into lethally irradiated congenic wild-type CD45.1 recipient mice followed by pIpC-induced *U2af1* deletion. Peripheral blood and bone marrow chimerism of *Mx1*-Cre, *U2af1*^flox/flox^, and *Mx1*-Cre/*U2af1*^wt/flox^ mice was compared to *Mx1*-Cre/*U2af1*^flox/flox^ mice. (B) Bulk peripheral blood chimerism (n=9-10). (C-D) Myeloid peripheral blood cell chimerism (n=9-10). (E-F) Lymphoid peripheral blood cell chimerism (n=9-10). (G) Bone marrow chimerism of mature hematopoietic cells (neutrophils, monocytes, B cells, and T cells) at ten months post-pIpC (n=8). (H) Bone marrow chimerism of stem and progenitor cells (KL, KLS, MPP, STHSC, LTHSC, SLAM) 10 months post-pIpC (n=8 for KL and KLS; n=3-4 for all other). All data are presented as mean +/− SD. *p<0.05, **p<0.01, ***p<0.001 by two-way (A-F) or one-way (G-H) ANOVA, with Tukey’s multiple comparison test. KL (Lineage^−^/cKit^+^/Sca1^−^), KLS (Lineage^−^/cKit^+^/Sca1^+^). KLS subpopulations: KLS subpopulations: SLAM (KLS/CD150^+^/CD48^−^), multipotent progenitors (MPP; KLS/CD34^+^/Flk2^+^), short-term stem cells (ST-HSC; KLS/CD34^+^/Flk2^−^), long-term stem cells (LT-HSC; KLS/CD34^−^/Flk2^−^).

### *U2af1* deletion increases expression of genes involved in splicing, unfolded protein response, and apoptosis

Adequate numbers of viable hematopoietic cells from adult mice following the conditional deletion of *U2af1* are difficult to obtain. Therefore, to examine the effects of reduced U2AF1 levels on expression of genes involved in hematopoiesis and hematopoietic cells survival, we performed RNA-sequencing on E14.5 hematopoietic myeloid progenitor cells (KL cells [CD45^+^/Lineage^−^/cKit^+^/Sca1^−^]). We first analyzed the expression of *U2af1* exon 2 in *U2af1*^flox/flox^ control cells, *Vav1*-Cre/*U2af1*^wt/flox^ cells (heterozygous KO), and *Vav1*-Cre/*U2af1*^flox/flox^ cells (homozygous KO). Exon 2 was flanked by loxP sites in our cells and should be deleted after Cre activation. *Vav1*-Cre/*U2af1*^wt/flox^ cells had the expected 50% reduction in exon 2 expression compared to *U2af1*^flox/flox^ control cells while *Vav1*-Cre/*U2af1*^flox/flox^ cells consistently had 70% reduced levels in exon 2 expression compared to *U2af1*^flox/flox^ control (i.e., ~30% residual levels compared to control cells). This residual expression is likely due to *U2af1* exon 2 expression from cells that escaped complete deletion of exon 2 (**Supplemental Figure 5A**).

Despite the incomplete deletion of *U2af1* in KO cells, unsupervised analysis of gene expression revealed that cells from the homozygous KO mice segregated away from heterozygous KO and control cells, which share a similar gene expression profile (**Figure Supplemental 5B**). Supervised analysis of gene expression in *Vav1*-Cre/*U2af1*^flox/flox^ compared to both *Vav1*-Cre/*U2af1*^wt/flox^ and control mice identified genes that were dysregulated in homozygous KO cells (**Figure 5A-C**). Notably, deletion of only one copy of *U2af1* (heterozygous KO) resulted in very few significant gene expression changes compared to the control cells (**Figure 5C**). Downregulated genes included enrichment in genes involved in myeloid differentiation, including *Csf1r* (**Figure 5D, Supplemental Figure 5D, Supplemental Table 2**). Genes involved in pre-mRNA splicing, response to unfolded protein, and apoptosis were significantly upregulated in homozygous KO cells compared to heterozygous KO and control cells (**Figure 5D, Supplemental Figure 5C, Supplemental Table 2**). Upregulation of genes involved in apoptosis was consistent with the previous report of U2AF1 knockdown in human erythroblasts (25) and our observed increase in apoptotic (annexin V^+^, viability-marker^−^) KL cells in the *Vav1*-Cre/*U2af1*^flox/flox^ mouse fetal liver cells compared to *Vav1*-Cre/*U2af1*^wt/flox^, *U2af1*^flox/flox^ and *Vav1*-Cre cells (**Supplemental Figure 2D**).

**Figure 5.**
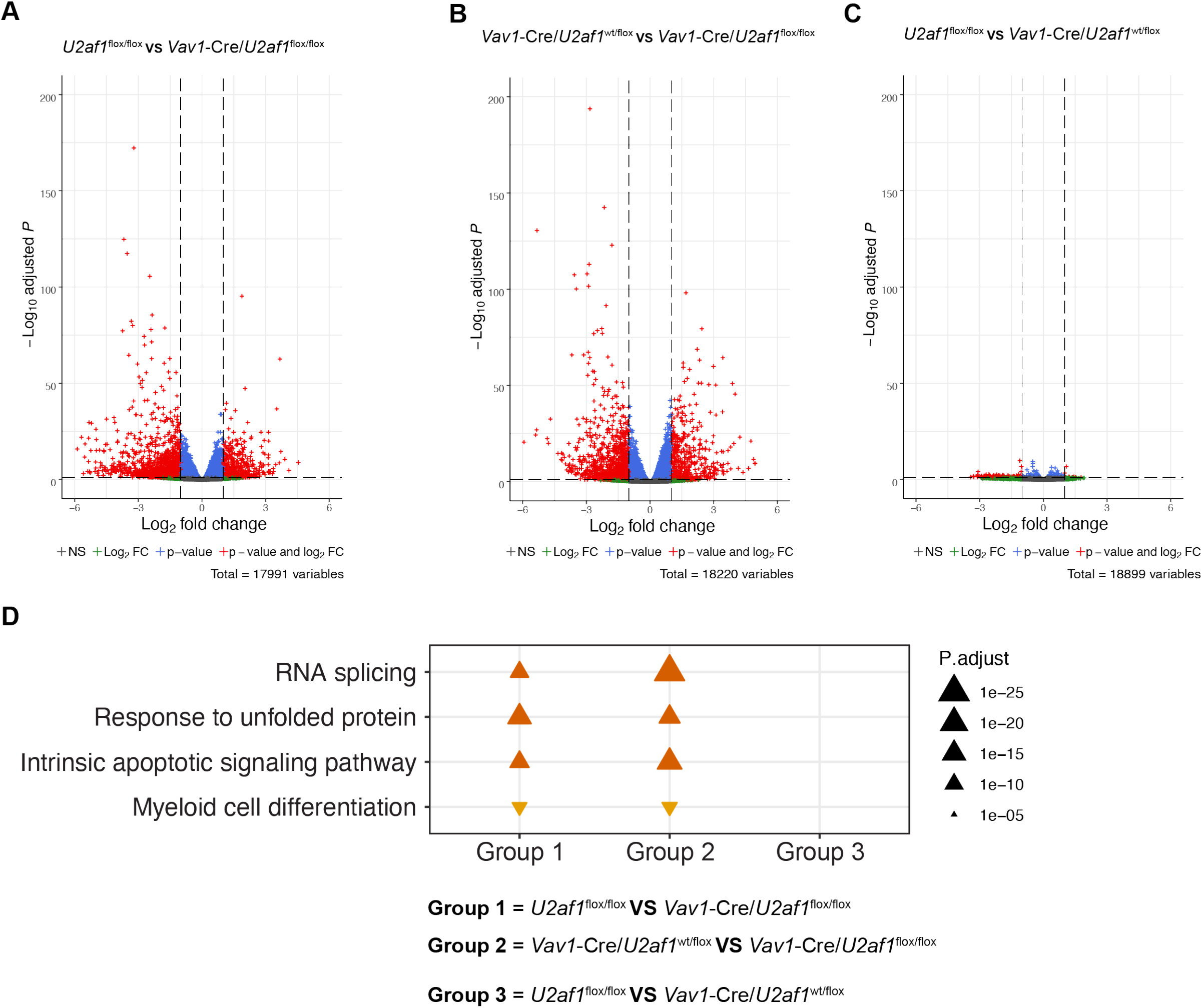
*U2af1* deletion dysregulates genes involved in splicing, unfolded protein response, and apoptosis pathways. Differential gene expression from RNA-seq of E14.5 hematopoietic progenitor cells (CD45^+^/Lineage^−^/cKit^+^/Sca1^−^). (A-C) Volcano plot comparing gene expression of (A) *Vav1*-Cre/*U2af1*^flox/flox^ compared to *U2af1*^flox/flox^ control, (B) *Vav1*-Cre/*U2af1*^flox/flox^ compared to *Vav1*-Cre/*U2af1*^wt/flox^, and (C) *Vav1*-Cre/*U2af1*^wt/flox^ compared to *U2af1*^flox/flox^ control. Red indicates genes that are significantly upregulated or downregulated at FDR<0.1 and fold-change>2. Blue are genes that are significantly regulated at FDR<0.1 and fold change<2. (D) Gene ontology (GO) enrichment analysis of the differentially regulated biological processes in *Vav1*-Cre/*U2af1*^flox/flox^ compared to *U2af1*^flox/flox^ control (group 1) and *Vav1*-Cre/*U2af1*^wt/flox^ (group 2), and *U2af1*^flox/flox^ control compared to *Vav1*-Cre/*U2af1*^wt/flox^ (group 3); there was no significantly dysregulated biological processes in Group 3. n=3 per genotype.

### 3’ consensus splice site sequence differs between *U2af1* KO and U2AF1 mutant cells

To analyze the effect of *U2af1* deletion on mRNA splicing, we used rMATS to identify and classify differentially spliced junctions. Unsupervised analysis of differentially spliced exons showed that the homozygous KO cells segregated away from heterozygous KO and control cells (**Figure 6A**). We identified 267 and 173 cassette exons that were skipped more in *Vav1*-Cre/*U2af1*^flox/flox^ cells compared to *U2af1*^flox/flox^ control and *Vav1*-Cre/*U2af1*^wt/flox^ cells, respectively, but only eight exons that were skipped more in *Vav1*-Cre/*U2af1*^wt/flox^ compared to the *U2af1*^flox/flox^ control cells (**Figure 6B**). Similar to the paucity of hematopoietic phenotypes and gene expression differences between heterozygous KO and control cells, RNA splicing is not dramatically altered in heterozygous KO cells.

**Figure 6.**
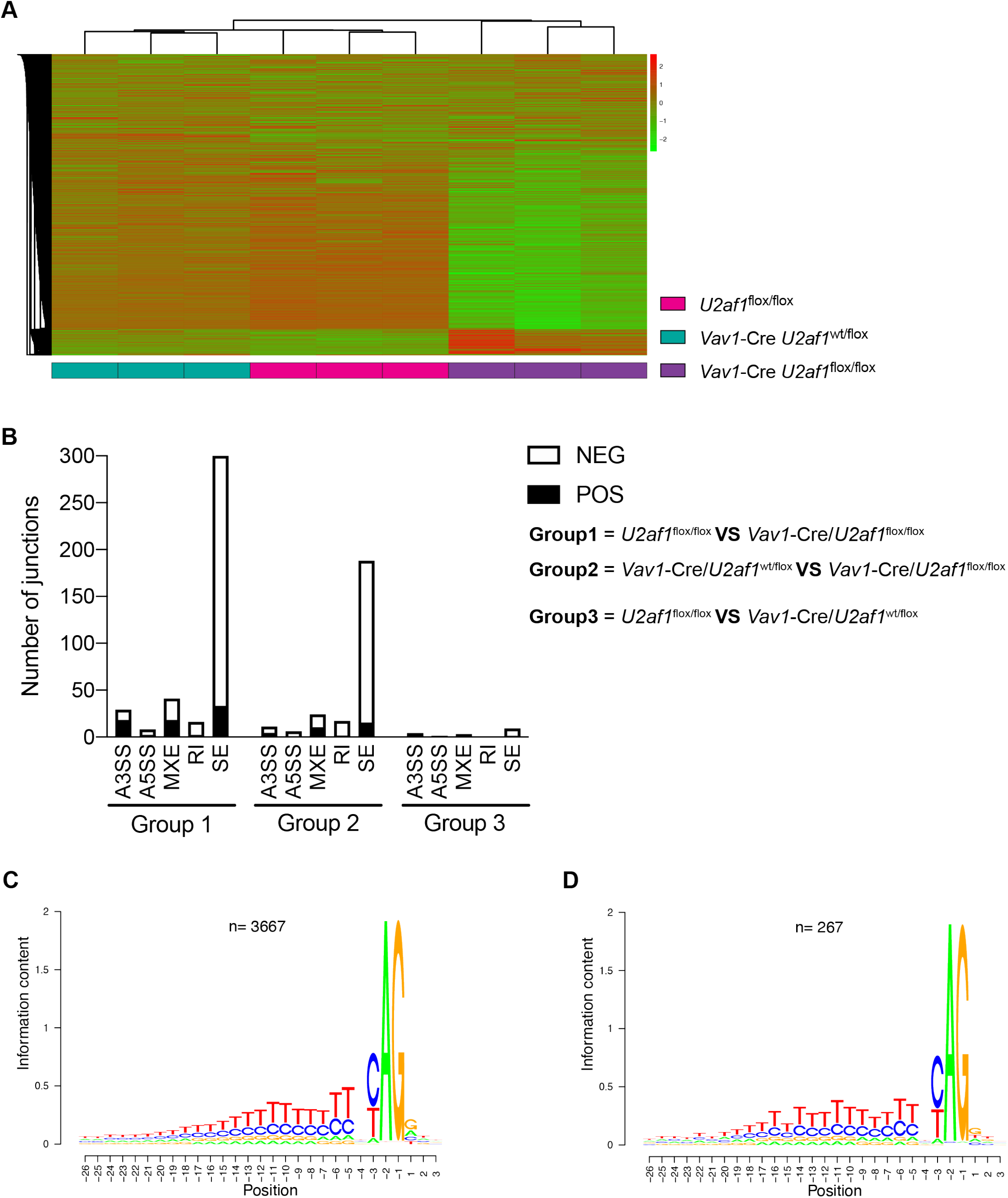
*U2af1* deletion induces alternative splicing but does not alter the specificity at the 3’ consensus splice site sequence of skipped exons. Differential splicing analysis of RNA-seq of E14.5 hematopoietic progenitor cells. (A) Unsupervised hierarchical clustering of the percent-spliced-in (PSI) of skipped exon junctions. (B) Distribution of alternative splicing events in *Vav1*-Cre/*U2af1*^flox/flox^ compared to *U2af1*^flox/flox^ control cells (left), *Vav1*-Cre/*U2af1*^flox/flox^ compared to *Vav1*-Cre/*U2af1*^wt/flox^ cells (center), and *Vav1*-Cre/*U2af1*^wt/flox^ compared to *U2af1*^flox/flox^ control cells (right). (C) The consensus sequence around the 3’ AG dinucleotide splice-acceptor site of cassette exons that are unchanged between *Vav1*-Cre/*U2af1*^flox/flox^ compared to *U2af1*^flox/flox^ cells (FDR>0.9, |ΔΨ|=0). (D) Intronic sequence context adjacent to the 3’ AG dinucleotide splice-acceptor site of splice sites skipped more often in *Vav1*-Cre/*U2af1*^flox/flox^ compared to *U2af1*^flox/flox^ (FDR<0.1, |ΔΨ|>0.1). SE (skipped exon), A5SS (alternative 5’ splice site), A3SS (alternative 3’ splice site), MXE (mutually exclusive exons), RI (retained intron). n=3 per genotype.

Further analysis of the skipped exons revealed that there was no sequence specificity at the 3’ splice site associated with the exons that were skipped more in the homozygous *U2af1* KO cells when compared to junctions that were not altered in KO cells (i.e., control splicing junctions) (**Figure 6C, D**). This lack of sequence specificity differs from what has been observed in mutant U2AF1-S34F/Y or U2AF1-Q157P/R expressing cells that have 3’ consensus splice site preferences that differ from the control consensus (26–29). Collectively, the splicing results suggest that *U2AF1* heterozygous KO cells do not phenocopy heterozygous hotspot mutations (coding for S34F/Y and Q157P/R), arguing against the mutations inducing loss-of-function properties, but likely confer gain- or neo-morphic functions (26–29). Pathway enrichment analysis using KEGG, Reactome, and Go-analysis of the differentially spliced genes found in homozygous *U2af1* KO cells revealed enrichment in genes involved in pre-mRNA splicing (**Supplemental Table 4**).

### Hematopoietic cells expressing mutant U2AF1(S34F) require expression of the residual wild-type allele for viability

We next examined whether mutant U2AF1(S34F)-expressing hematopoietic cells require the expression of the residual wild-type allele for survival and directly tested whether *U2AF1* is a haplo-essential gene *in vivo*. We crossed the previously published *U2af1* heterozygous knock-in mouse [*U2af1*^wt/S34F^ KI] (29) expressing the serine to phenylalanine substitution at amino acid 34 (S34F) from its endogenous locus to the *U2af1* KO mouse to generate the following genotypes: *Mx1*-Cre*, Mx1*-Cre/*U2af1*^wt/flox^, *Mx1*-Cre/*U2af1*^flox/flox^, *Mx1*-Cre/*U2af1*^wt/S34F^ [heterozygous *U2af1*^wt/S34F^ KI], and *Mx1*-Cre/*U2af1*^flox/S34F^ [hemizygous *U2af1*^−/S34F^ KI]. We then performed a competitive repopulation transplantation assay (**Figure 7A**). We monitored the neutrophil chimerism because neutrophils were rapidly depleted within two weeks in our homozygous *U2af1* KO model, providing a fast and sensitive measure of cell survival. As expected, at two and four weeks post-pIpC-induced *U2af1* deletion, *Mx1*-Cre only control and heterozygous *U2af1* KO neutrophil chimerism were unchanged, while the homozygous *U2af1* KO neutrophils were depleted quickly, and the heterozygous *U2af1*^wt/S34F^ KI neutrophils had a previously reported disadvantage (29) (**Figure 7B**). At two and four weeks post-pIpC, the hemizygous *U2af1*^−/S34F^ KI cells were not viable (**Figure 7B**), indicating that mutant U2AF1(S34F) expression cannot rescue the *U2af1* KO-induced lethality, and mutant U2AF1(S34F) hematopoietic cells require the expression of the wild-type U2AF1 for cell survival *in vivo*, consistent with *U2AF1* being a haplo-essential gene.

**Figure 7.**
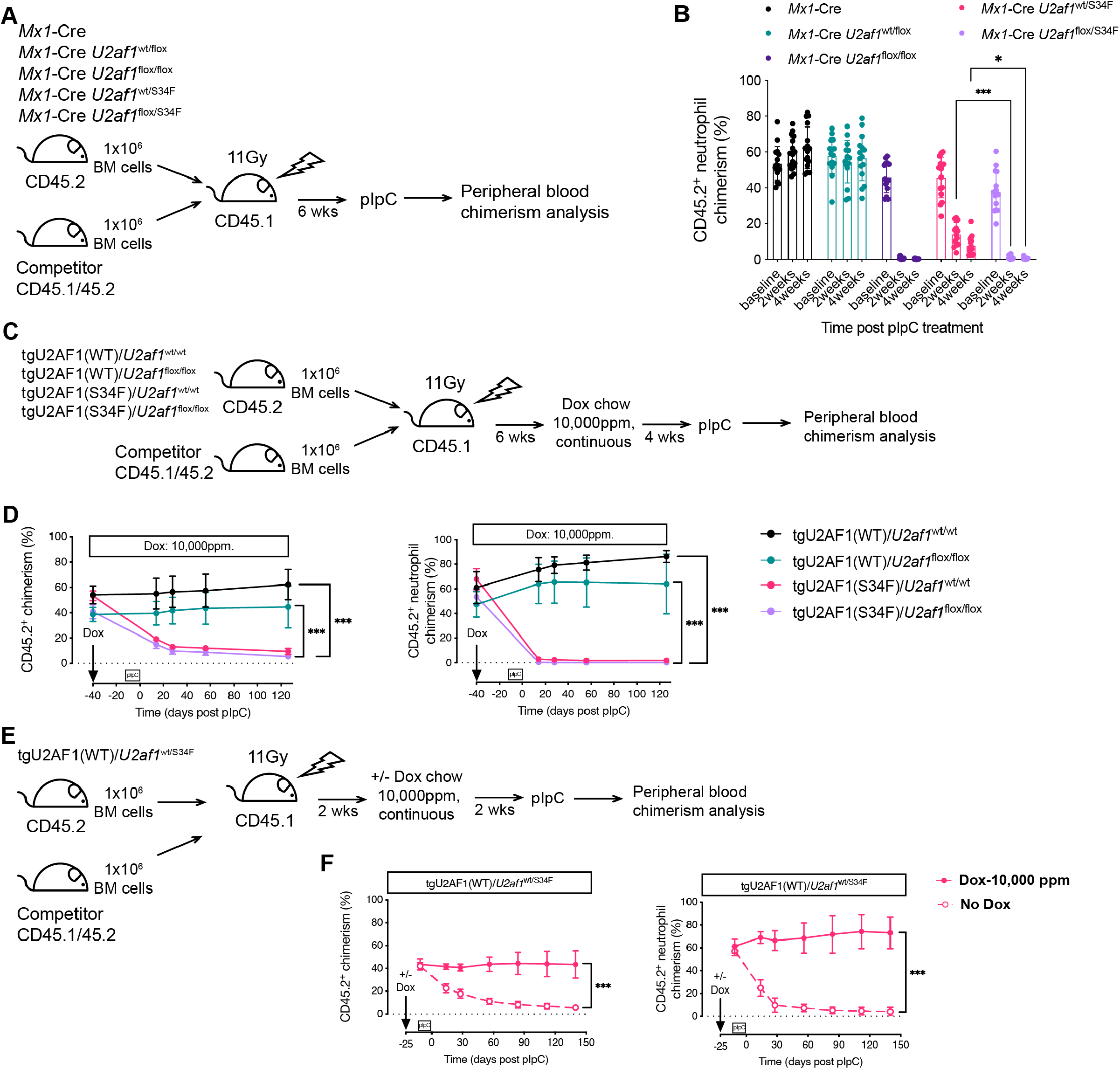
Survival of mutant U2AF1(S34F) hematopoietic cells is dependent on the expression of the residual wild-type allele and the ratio of U2AF1(WT:S34F) expression. (A) Experimental design of a competitive transplant of whole bone marrow cells from CD45.2 *Mx1*-Cre, *Mx1*-Cre/*U2af1*^wt/flox^, *Mx1*-Cre/*U2af1*^flox/flox^, *Mx1*-Cre/*U2af1*^wt/S34F^, or *Mx1*-Cre/*U2af1*^flox/S34F^ mice mixed at a 1:1 ratio with congenic wild-type CD45.1/45.2 competitor cells followed by transplantation into lethally irradiated congenic wild-type CD45.1 recipient mice. *U2af1* deletion was induced by pIpC, and analysis of the peripheral blood chimerism was performed. (B) Peripheral blood neutrophil chimerism of *Mx1*-Cre, *Mx1*-Cre/*U2af1*^wt/flox^, *Mx1*-Cre/*U2af1*^flox/flox^, *Mx1*-Cre/*U2af1*^wt/S34F^, and *Mx1*-Cre/*U2af1*^flox/S34F^ mice (n=14-16). (C) Experimental design of a competitive transplant of whole bone marrow cells from CD45.2 tgU2AF1(WT)/*U2af1*^wt/wt^, tgU2AF1(WT)/*U2af1*^flox/flox^, tgU2AF1(S34F)/*U2af1*^wt/wt^, or tgU2AF1(S34F)/*U2af1*^flox/flox^ mice. All the donor test mice have both *Mx1*-Cre and rtTA. Transgenic U2AF1(WT) and U2AF1(S34F) were induced by 10,000 ppm doxycycline chow (dox chow) followed by pIpC-induced *U2af1* deletion after four weeks, and analysis of the peripheral blood chimerism was performed. (D) Overall peripheral blood and neutrophil chimerism in tgU2AF1(WT)/*U2af1*^wt/wt^, tgU2AF1(WT)/*U2af1*^flox/flox^, tgU2AF1(S34F)/*U2af1*^wt/wt^, and tgU2AF1(S34F)/*U2af1*^flox/flox^ mice after induction of transgenic U2AF1(WT) or U2AF1(S34F) by 10,000 ppm dox chow and *U2af1* deletion induced by pIpC Cre-activation (n=7-8). (E) Experimental design of a competitive transplant of whole bone marrow cells from CD45.2 *Mx1*-Cre/rtTA/tgU2AF1(WT)/*U2af1*^wt/S34F^ (tgU2AF1[WT]/*U2af1*^wt/S34F^) mice. Transgenic U2AF1(WT) was induced by 10,000 ppm dox chow two weeks post-transplant followed by pIpC-induced *U2af1* deletion after two weeks of dox chow and analysis of peripheral blood chimerism was performed. (F) Overall peripheral blood and neutrophil chimerism of tgU2AF1(WT)/*U2af1*^wt/S34F^ with or without dox chow treatment (n=9-10). All data are presented as mean +/− SD. *p<0.05, ***p<0.001 by two-way ANOVA with Sidak’s (B, F) or Tukey’s (D) multiple comparison test.

While hemizygous expression of mutant U2AF1 in cells is lethal (expression of only one mutant U2AF1 allele), we next asked whether increasing mutant U2AF1(S34F) expression to a higher level could rescue the *U2af1* KO-induced lethality using a complementation test. We used our previously characterized doxycycline-inducible transgenic mouse models that express either U2AF1(S34F) or U2AF1(WT) (28). The transgenic mice harbor the reverse tetracycline-controlled transactivator (rtTA) inserted into the *Rosa26* locus, allowing for transgenic U2AF1 expression following doxycycline exposure. We first intercrossed tgU2AF1(WT) or tgU2AF1(S34F) mice with *Mx1*-Cre/*U2af1*^flox/flox^ mice to generate tgU2AF1(WT)/*Mx1*-Cre/*U2af1*^flox/flox^ or tgU2AF1(S34F)/*Mx1*-Cre/*U2af1*^flox/flox^ mice and harvested donor test hematopoietic cells to use in a competitive repopulation transplant assay. Donor test cells were mixed with an equal number of wild-type congenic competitor cells and transplanted into lethally-irradiated congenic recipient mice. Following complete engraftment, recipient mice were fed 10,000 ppm doxycycline chow to induce expression of tgU2AF1(WT) or tgU2AF1(S34F) (**Figure 7C**). After deleting endogenous *U2af1*, we observed that overexpression of tgU2AF1(WT), but not mutant tgU2AF1(S34F), was able to rescue the homozygous *U2af1* KO-reduced peripheral blood chimerism, including neutrophils, B cells, and T cells chimerism (**Figure 7D, Supplemental Figure 6B, C**). The results indicate that mutant U2AF1 lacks critical wild-type functions that are necessary to rescue the lethality observed in *U2af1* KO cells, even when expressed at higher levels.

### The ratio of U2AF1 wild-type to mutant S34F expression determines the phenotype of mutant cells

Previous *in vitro* studies using a human lung epithelial cell line genetically modified to express mutant U2AF1 showed that the ratio of U2AF1 wild-type to mutant S34F expression (U2AF1[WT:S34F]) determines the phenotype of mutant expressing cells (17). To test whether the ratio of U2AF1(WT:S34F) expression affects hematopoietic phenotypes *in vivo,* we used two complementary approaches (e.g., overexpress U2AF1[WT] to increase the U2AF1[WT:S34F] expression ratio or overexpress mutant U2AF1[S34F] to decrease the ratio). We first intercrossed tgU2AF1(WT) mice with *Mx1*-Cre/*U2af1*^wt/S34F^ mice to generate tgU2AF1(WT)/*Mx1*-Cre/*U2af1*^wt/S34F^ donor mice. We harvested test hematopoietic cells from donor mice, mixed them with equal numbers of wild-type congenic competitor cells, and transplanted them into a lethally irradiated congenic recipient mice for a competitive bone marrow transplant (**Figure 7E**). We fed recipient mice 10,000 ppm doxycycline chow to overexpress U2AF1(WT) and increase the ratio of wild-type to mutant U2AF1 expression in test donor cells. We observed that the competitive disadvantage of neutrophils and other lineages in the heterozygous *U2af1*^wt/S34F^ KI mice could be fully rescued by overexpressing tgU2AF1(WT) (**Figure 7F, Supplemental Figure 6F, G**).

Next, we tested whether decreasing the ratio of wild-type to mutant U2AF1 expression by increasing the U2AF1(S34F) expression level would worsen the competitive disadvantage of mutant cells, as predicted. We harvested donor test cells from tgU2AF1[S34F]/*U2af1*^wt/wt^ mice and mixed them with equal numbers of congenic competitor donor bone marrow cells before injection into lethally irradiated congenic recipient mice (28). When we increased the dose of doxycycline from 625 ppm to 10,000 ppm to increase the expression level of U2AF1(S34F), the competitive disadvantage of mutant U2AF1(S34F) cells worsened (**Supplemental Figure 6D**). Collectively, these data indicate that reducing the ratio of wild-type to mutant U2AF1 expression, by either reducing wild-type expression or increasing mutant expression, can induce lethality of hematopoietic cells *in vivo*. This indicates that modulating the ratio of wild-type to mutant U2AF1 expression could be an approach to target mutant cells.

### Hematopoietic cells expressing mutant U2AF1(S34F) are more sensitive than non-mutant cells to reduced levels of wild-type U2AF1

Next, we assessed if the mutant U2AF1(S34F)-expressing cells are more sensitive than non-mutant cells to reduced but not absent levels of wild-type U2AF1, rather than complete loss of wild-type U2AF1 achieved in hemizygous mutant cells. We generated tgU2AF1(WT)/*U2af1*^flox/flox^, and tgU2AF1(WT)/*U2af1*^flox/S34F^ mice, both also expressing *Mx1*-Cre and rtTA, allowing us to delete endogenous wild-type *U2af1* and control the level of U2AF1(WT) using various doxycycline doses. We performed a bone marrow competitive repopulation transplantation assay (**Figure 8A**). Two weeks post-transplantation, we induced tgU2AF1(WT) expression using 10,000 ppm doxycycline chow followed by pIpC-induced deletion of the endogenous mouse wild-type *U2af1* allele and expression of the mutant allele. As expected, with 10,000 ppm doxycycline chow (used to express transgenic U2AF1[WT]), both *U2af1* homozygous KO and *U2af1*^−/S34F^ hemizygous KI neutrophils were viable. However, as the dose of transgenic U2AF1(WT) was dropped by decreasing the doxycycline dose from 10,000 ppm to 2500 ppm or 1250 ppm, the U2AF1^S34F^-expressing neutrophils were more vulnerable to the reduced doses (corresponding to lower levels of tgU2AF1[WT]) compared to neutrophils expressing only the tgU2AF1(WT) (**Figure 8B, C**). These results suggest there is a wild-type U2AF1 expression threshold below which U2AF1 mutant-expressing cells cannot survive, but wild-type cells can. This threshold may create a vulnerability in mutant cells that could be exploited therapeutically to preferentially kill mutant cancer cells.

**Figure 8.**
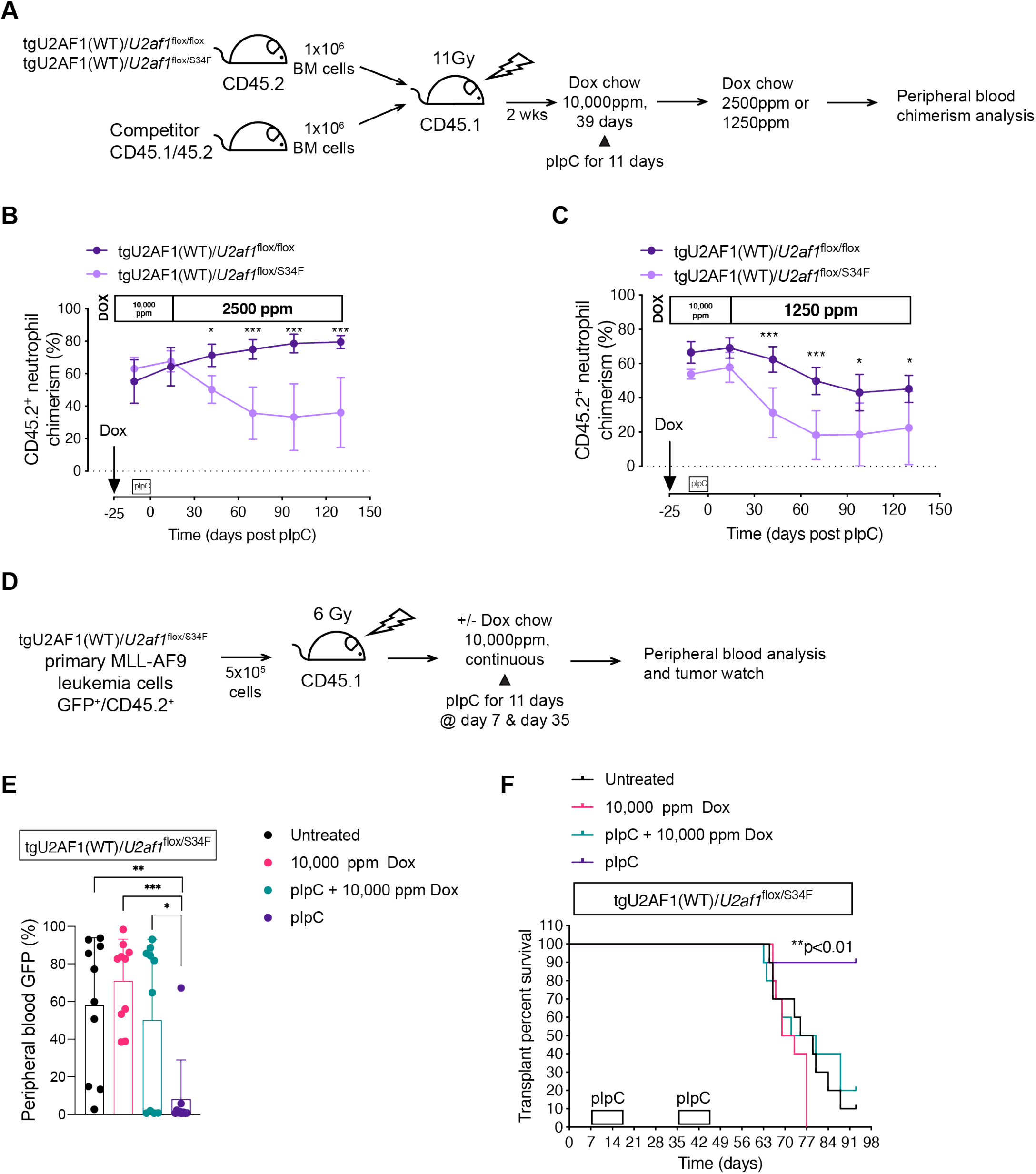
Hematopoietic cancer cells expressing mutant U2AF1(S34F) are sensitive to decreased levels of the wild-type U2AF1 expression. (A) Experimental design of a competitive transplant of test whole bone marrow cells from CD45.2 tgU2AF1(WT)/*U2af1*^flox/flox^ or tgU2AF1(WT)/*U2af1*^flox/S34F^ mice mixed at a 1:1 ratio with congenic wild-type CD45.1/45.2 competitor cells followed by transplantation into lethally irradiated congenic wild-type CD45.1 recipient mice. All the donor test mice have both *Mx1*-Cre and rtTA. Transgenic U2AF1(WT) expression was induced by 10,000 ppm doxycycline chow (dox chow) two weeks post-transplant followed by pIpC induction of *U2af1* deletion two weeks later. Two weeks post-pIpC, the dox dose was reduced to 2500 ppm or 1250 ppm, and analysis of the peripheral blood chimerism was performed. (B) Peripheral blood neutrophil chimerism of tgU2AF1(WT)/*U2af1*^flox/flox^ and tgU2AF1(WT)/*U2af1*^flox/S34F^ after induction of transgenic U2AF1(WT) by 2500 ppm dox chow (n=5). (C) Peripheral blood neutrophil chimerism of tgU2AF1(WT) *U2af1*^flox/flox^ and tgU2AF1(WT)*/U2af1*^flox/S34F^ after induction of transgenic U2AF1(WT) by 1250 ppm dox chow (n=5). (D) Experimental design of transplantation of tgU2AF1(WT)/*U2af1*^flox/S34F^ MLL-AF9 AML tumor cells (GFP^+^/ CD45.2^+^) isolated from the spleen of primary mice into sub-lethally irradiated secondary recipients. Secondary recipients were treated with or without 10,000 ppm doxycycline chow followed by pIpC induction and analysis of the peripheral blood analysis and tumor watch. (E) GFP^+^ MLL-AF9 AML cells chimerism up to 21 days post-second pIpC dose (n=10). (F) Kaplan-Meier survival curves up to 93 days post-transplant (n=10). All data are presented as mean +/− SD. *p<0.05, **p<0.01, ***p<0.001, by two-way (B, C) or one-way (E, F) ANOVA with Tukey’s multiple comparison test.

It is possible that the viability of fully transformed U2AF1 mutant cells is not dependent on U2AF1 wild-type expression. To test this, we generated primary MLL-AF9 AML tumors in mice by transducing tgU2AF1(WT)/*U2af1*^flox/S34F^ mouse hematopoietic cells with MLL-AF9 retrovirus before exposure of doxycycline or pIpC. We then transplanted the MLL-AF9 AML tumor cells isolated from primary mice into sub-lethally irradiated secondary recipients (**Figure 8D**). Mice were then treated with or without 10,000 ppm doxycycline chow to induce transgenic U2AF1(WT) expression prior to vehicle or pIpC induction (to delete the endogenous wild-type *U2af1* and express mutant U2AF1[S34F]). This approach allowed us to control the level of U2AF1(WT) expression using doxycycline. Mice that received only pIpC (i.e., creating *U2af1*^−/S34F^ hemizygous AML cells in recipients) had significantly lower tumor burden in the peripheral blood and increased survival compared to all other mice (e.g., mice harboring U2AF1 mutant AML that express wild-type U2AF1) and mice that received both 10,000 ppm doxycycline and pIpC (i.e., hemizygous AML cells expressing exogenous U2AF1[WT]) (**Figure 8E, F; Supplemental Figure 7E-G**). These results suggest that fully transformed primary U2AF1 mutant AML cells are also sensitive to the ratio of U2AF1 wild-type to mutant expression for survival.

## Discussion

In MDS and other cancers, *U2AF1* and other spliceosome gene mutations are heterozygous, and the residual wild-type allele is expressed. This observation suggests that the wild-type allele is required for the survival of mutant U2AF1 expressing cells. Using *U2af1* conditional knock-out and S34F knock-in mice, our results demonstrate that *U2AF1* is a haplo-essential cancer gene. Our data also shows that modulation of the ratio of U2AF1 wild-type:S34F mutant expression is critical for hematopoietic cell survival *in vivo*, including leukemia cells. We show that lowering U2AF1 wild-type or increasing mutant U2AF1(S34F) expression in mutant cells reduces cell viability, implicating that modulation of the ratio of wild-type:mutant U2AF1 expression may be a potential therapeutic strategy in cancer.

Our data also provide evidence that *U2AF1* mutations are not simply creating a haploinsufficient state in mutant cells because heterozygous *U2af1* KO mice are normal. In addition, heterozygous KO progenitor cells have very few changes in gene expression or splicing alterations compared to control cells. Additional findings, including that deletion of *U2af1* does not change the sequence specificity at the 3’ splice site of alternatively spliced junctions, unlike mutant U2AF1, (26–30) suggest that mutant U2AF1 confers a change-of-function rather than loss-of-function. Accumulating evidence indicates that hotspot mutations in the splicing factors *SF3B1* and *SRSF2* also cause change-in-function (i.e., neomorphic functional change) rather than loss-of-function properties (31–37).

*U2AF1* was recently identified as a haplo-essential cancer gene based on computational analysis of sequencing data. Unlike other oncogenic driver gene mutations that undergo allelic imbalance (e.g., copy number alteration [amplification or deletion] or uniparental disomy) to increase the mutant allele dosage over the wild-type allele, spliceosome gene mutations undergo negative selection to preserve and skew towards the wild-type allele when an allelic imbalance occurs in a tumor (12). Our results show that hemizygous (i.e., one mutant and one deleted allele) hematopoietic cells are not viable and have a strong competitive disadvantage in competitive repopulation assays compared to wild-type cells, confirming that the wild-type allele is required for viability, and consistent with *U2AF1* being a haplo-essential cancer gene. Our results are also consistent with previous *in vitro* results using human lung epithelial cell line expressing mutant U2AF1 as well as *in vivo* data using a mouse model of mutant SRSF2, both showing that hemizygous cells are not viable (14, 17, 38, 39). Therefore, a common feature shared by recurrent *SRSF2, U2AF1,* or *SF3B1* heterozygous mutations is likely their sensitivity to loss or reduced expression of the residual wild-type allele.

The dependence of cell viability on the ratio of the wild-type to mutant allele expression was further verified using several approaches. In addition to reducing mutant cell viability by deleting the *U2af1* wild-type allele, overexpressing mutant U2AF1 also reduced cell viability. Overexpressing mutant U2AF1(S34F) using a doxycycline-inducible mouse model resulted in a dose-dependent reduction in cell viability *in vivo*. Finally, overexpression of the wild-type allele above the endogenous heterozygous mutant levels rescued the hematopoietic cell effects induced by mutant U2AF1 *in vivo*. Collectively, these results consistently show that the ratio of the wild-type to mutant expression drives hematopoietic phenotypes *in vivo* (17).

The dependence of cell viability on the expression of the wild-type *U2AF1* allele in mutant U2AF1(S34F) expressing cells was also observed using primary MLL-AF9 mouse AML samples whose growth was significantly hindered in the hemizygous state. Overall, these results suggest that targeting (i.e., reducing) the expression of the wild-type U2AF1 allele in mutant-expressing cancer cells could be a therapeutic strategy. While drugs that directly target wild-type U2AF1 have not been reported, antisense oligonucleotides (ASO) could be used to directly reduce (not completely) the expression of wild-type but not the mutant U2AF1. Specificity has been achieved at a single nucleotide level for ASOs targeting another gene harboring a mutation (40).

Recent progress in targeting spliceosome mutant cancer cells has expanded. The preclinical efficacy of splicing modulators targeting SF3B1 have been reported in cells bearing mutations in *U2AF1, SRSF2,* and *SF3B1*. These studies have shown that mutant cells are more sensitive to pharmacological inhibition of SF3B1 relative to the wild-type cells (13, 14, 16). In addition, spliceosome mutant cells are more sensitive than wild-type cells to inhibition of ATR, a regulator of R-loops (41). These approaches are now being tested in spliceosome mutant patients with myeloid neoplasms, including a modulator of SF3B1, H3B-8800 (NCT02841540), and an ATR inhibitor (NCT03770429). As our understanding of the mechanisms underlying the vulnerability of spliceosome mutant cells continues to expand, the introduction of novel therapies to eradicate spliceosome mutant cells could improve the outcomes for MDS patients with spliceosome gene mutations.

## Methods

### Generating *U2af1* KO mouse

Details of generating the *U2af1* KO are in the supplemental methods. To create the conditional *U2af1* KO mouse, exon 2 was flanked with loxP sites in targeting vector. Targeted ES cell clones were injected into C57BL/6 blastocysts and implanted to create chimeras. Chimeras were bred to wild-type C57BL/6 mice to generate a mouse with germline transmission of the *U2af1* KO for use in this study.

### Bone marrow transplantation assays

Bone marrow was harvested from donor mice femurs, tibias, and iliac crests, then resuspended in FACS buffer (PBS without Ca^2+^ and Mg^2+^, 2% FBS, and 1 mM EDTA). The cells were filtered through a 70μm cell strainer (Corning) then counted using a Cellometer Auto 2000 cell viability counter (Nexcelom Bioscience). For non-competitive bone marrow transplantation, CD45.2 donor mice bone marrow was resuspended in Hanks’ Balanced Salt Solution (HBSS), and then 2×10^6^ cells were transplanted by retroorbital injection into lethally irradiated (11 Gy) congenic wild-type CD45.1 recipient mice. For competitive repopulation assay, whole bone marrow cells from CD45.2 mice were mixed at a 1:1 ratio with congenic wild-type CD45.1/45.2 competitor cells and resuspended HBSS followed by transplantation of 2×10^6^ cells by retroorbital injection into lethally irradiated (11 Gy) congenic wild-type CD45.1 recipient mice. Following pIpC-induced *U2af1* deletion, peripheral blood chimerism was monitored by flow cytometry.

### Bone marrow cell viral transduction and transplantation

Bone marrow cells were isolated as described above then resuspended in stem cell culture media composed of Stempro-34 medium (Gibco), StemPro-34 nutrient supplement, Pen-Strep (Thermo Fisher), GlutaMAX™ (Thermo Fisher), and murine cytokines (SCF [100 ng/mL], TPO [10 ng/mL], Flt3 [50 ng/mL] and IL3 [6 ng/mL]). For the transduction of cells with MLL-AF9 virus, polybrene was added, followed by spinfection with lentivirus preparation at (1500 rpm) at 35° C for 90 minutes. The cells were then cultured at 37° C overnight and spinfected again. After another overnight culture, the cells were transplanted into lethally irradiated recipients 24 hours post-transduction by tail vein injection. The harvested primary tumors were used for subsequent secondary transplantation.

### Peripheral blood analysis

Peripheral blood counts were assessed using automated complete blood count (Hemavet 950, Drew Scientific).

### Cell Staining and Flow Cytometry

The red blood cells in the bone marrow, peripheral blood, spleen, and fetal liver were depleted using ammonium chloride potassium bicarbonate (ACK; 150 mM NH_4_Cl, 10 mM KHCO_3_, 0.1 mM Na_2_EDTA) lysis buffer followed by resuspension in FACS buffer for staining. Flow cytometry of mature lineage hematopoietic cells was performed using antibodies targeting cell surface receptors CD115, Gr-1, B220, and CD3e. Identification of the donor-derived cells (test or competitor) and the recipient cells was achieved using anti-CD45.1 and anti-CD45.2 antibodies. CD45.2 was also used to mark fetal liver hematopoietic cells. For hematopoietic stem and progenitor staining, a cocktail of antibodies against Gr-1, CD3e, B220, Ter119, and CD41 were used to exclude the mature lineage cells. Following the lineage mature cell exclusion, antibodies against cKit, Sca-1, CD34, FLT3, CD150, CD48, and FcγR were used to mark the stem and progenitor cells. All staining antibodies and their clone identities are listed in **Supplemental Table 5**. FACS analysis was performed on ZE5 Cell Analyzer (Bio-Rad) or Gallios (Beckman Coulter) flow cytometers. Cell sorting was done on iCyt Synergy sorter (Sony). Data analysis was performed using FloJo software (FlowJo LLC).

### Methylcellulose colony-forming assay

After ACK lysis of red blood cells, E14.5 fetal liver cells were resuspended in FACS buffer, and the cells filtered through a 70μm cell strainer. From each genotype, 25,000 cells were plated in duplicate in MethoCult GF M3434 medium (Stem Cell Technologies) and cultured at 37°C with 5% CO_2_. The number of colony-forming units (CFUs) was counted eight days later under an inverted microscope (Nikon TMS).

### Histological analysis

The tibia was fixed in 10% neutral buffered formalin (EKI, Joliet, IL) for a minimum of 24 hours at room temperature. The bones were decalcified in an HCl-EDTA decalcifier (EKI, Joliet, IL) for approximately 1 hour. The bones were processed for paraffin embedding using standard techniques, then embedded in paraffin and sectioned at 5 μm, followed by H & E staining. Images were captured using Leica Application Suite (Leica Microsystems GmbH, Wetzlar, Germany).

### RNA-Sequencing and analysis

RNA was isolated from sorted E14.5 fetal liver hematopoietic progenitor cells (KL; CD45^+^/Lineage^−^/cKit^+^/Sca1^−^) using the NucleoSpin RNA Plus XS kit (Macherey-Nagel). The RNA-sequencing libraries were prepared using KAPA RNA Hyper Prep Kit with RiboErase. After amplification, ~300 bp fragments were sequenced on Illumina NovaSeq 6000. The reads were then mapped using HISAT2 (version 2.1.0) (42) against the GRCm38 version of the mouse genome from Ensembl consortium. Gene level counts were processed to exclude any secondary or unmapped reads using the SAMtools and counts were generated using HTseq (version 0.11.0) (43). Differentially expressed genes were identified using DESeq2 (44). Additional filtering was applied to require genes to have five reads in at least half the samples, and a False Discovery Rate (FDR) <0.1. TPM values were calculated using StringTie (version 1.3.3) (45). Splicing event classification was performed using rMATS (version 4.0.2) (46). Gene enrichment analysis was performed using clusterProfiler (version 3.10.1) (47). Pathway enrichment analysis was performed using fgsea (version 1.8.0) against GO, Reactome, and MSIGDB gene sets (FDR < 0.1) (48).

## Data and code availability

The RNA-seq data generated in this study are available at NCBI’s Gene Expression Omnibus (GEO) repository with an accession number GSE152213.

## Statistics

Analysis of variance (ANOVA) followed by Tukey’s, Sidak’s, or Dunnett’s multiple comparison test was used for statistical comparison. Chi-Square test was used to test the difference between observed and the expected frequencies from different genotypes in *Vav1*-Cre *U2af1* KO breeding. Survival Kaplan-Meier curves were analyzed by the Mantel-Cox log-rank test. All data are presented as mean +/− SD. All statistical data analysis and graph plotting was done on GraphPad Prism 8 software (GraphPad Software), unless otherwise stated.

## Study approval

All mouse experiments were performed per institutional guidelines for care and use of laboratory animals and approved by the Washington University Animal Studies Committee.

## Author contributions

BAW and MJW designed the study. BAW, AH, MOA, MN, SG, JB, JS, TA, CLS, and DLF performed experiments. BAW, AH, SNS, CAM, and AK analyzed data. BAW, AH, SNS, TAG, and MJW interpreted the data, wrote and edited the manuscript. All others reviewed and approved the manuscript.

## Acknowledgments

This study was supported by grants from the Edward P. Evans Foundation, Lottie Caroline Hardy Trust, Taub Foundation, Siteman Investment Program - Barnard Trust, Siteman Investment Program - The Foundation for Barnes-Jewish Hospital Cancer Frontier Fund, and the Specialized Program of Research Excellence in AML (P50CA171963, to Dr. TAG and Dr. MJW). A grant from the Department of Defense (CA150844, to Dr. BAW). Support by the National Cancer Institute of the National Institutes of Health under Award Number K12 CA167540. We thank the Alvin J. Siteman Cancer Center at Washington University School of Medicine and Barnes-Jewish Hospital in St. Louis, MO., for the use of the Siteman Flow Cytometry. The Siteman Cancer Center is supported in part by an NCI Cancer Center Support Grant #P30 CA091842. Technical assistance was provided by the DCM Research Animal Diagnostic Laboratory. The authors thank Dr. Sanghyun Kim, Julie Ritchey, and Gayla Hadwiger for technical assistance. We thank Dr. Harold Varmus for providing the U2AF1(S34F) KI mouse and Drs. Tim Ley, Dan Link, Laura Schuettpelz, and Eugene Oltz for helpful scientific discussions.

## Notes

**Conflict of interest statement:** The authors have declared that no conflict of interest exists

### Competing Interest Statement

The authors have declared no competing interest.

