## Supplemental Materials for "*U2AF1* is a haplo-essential gene required for cancer cell survival"

**Authors and affiliations**

Brian A. Wadugu,[1] Amanda Heard,[1] Sridhar N. Srivatsan,[1] Michael O. Alberti,[2] Matthew Ndonwi,[1] Sarah Grieb,[1] Joseph Bradley,[1] Jin Shao,[1] Tanzir Ahmed,[1] Cara L. Shirai,[1] Ajay Khanna, [1] Dennis L. Fei, [3,4] Christopher A. Miller, [1] Timothy A. Graubert,[5] Matthew J. Walter[1]

[1]Division of Oncology, Department of Medicine, [2]Department of Pathology and Immunology, Washington University, St. Louis, MO 63110, USA

[3]Department of Medicine, Meyer Cancer Center, Weill Cornell Medicine, NY 10065, USA

[4]Cancer Biology Section, Cancer Genetics Branch, National Human Genome Research Institute, Bethesda, MD 20892, USA

[5]Massachusetts General Hospital Cancer Center, Harvard Medical School, Boston, MA 02114, USA.

### Supplemental methods

#### Animal models

All mice used in this experiment were of the C57BL/6 background. The *U2af1* KO mouse was generated by Ingenious Targeting Laboratory (2200 Smithtown Avenue, Ronkonkoma, NY 11779) using standard gene targeting techniques. To create the conditional *U2af1* KO mouse, exon 2 was flanked with loxP sites in targeting vector. The 5' single loxP site was engineered 443 bp upstream of exon 2 outside a potential regulatory region. The loxP/FRT-flanked Neo cassette, which served as the 3' loxP site after FLP-mediated Neo deletion, was placed 217 bp downstream of exon 2. The long homology arm extended about 6 kb 5' to the loxP cassette, and the short homology arm was about 2.1 kb 3' to the Neo cassette. The targeting vector was linearized and then transfected by electroporation of FLP C57BL/6 embryonic stem (ES) cells. After selection with G418 antibiotic, surviving clones were expanded for PCR analysis to identify recombinant ES cell clones. The Neo cassette in the targeting vector was removed during ES cell clone expansion. Secondary confirmation of positive clones identified by PCR was performed by Southern Blotting analysis. DNA was digested with restriction enzymes AvrII and MfeI to confirm the integration of targeting vector. The digested DNA was separated by gel electrophoresis on a 0.8% agarose gel. After transfer to a nylon membrane, the digested DNA was hybridized with a probe targeted against the target region. DNA from the C57Bl/6 mouse strain was used as wild type control. Targeted ES cells were confirmed to have a normal karyotype. The ES cell clones were injected into C57BL/6 blastocysts and implanted to create chimeras. Chimeras were bred to wild-type C57BL/6 mice, and germline transmission of the *U2af1* KO allele was verified in two mice obtained from one targeted ES cell clone by PCR and whole-genome sequencing. The mice were genotyped using the 5'

GAGGGTGATGGACCTACCCA 3' forward, and 5' GAGAAGAAAGGGGCGGCAAA 3' reverse primers (Targeted=295 bp, wild-type=112 bp). Mice carrying the *U2af1* floxed allele were crossed to the existing C57BL/6 Cre-expressing lines, including Cre-*ERT2*, *Mx1*-Cre, or *Vav1*-Cre as well as existing C57BL/6 doxycycline-inducible transgenic U2AF1(WT), transgenic U2AF1(S34F), or *U2af1*(S34F) KI mice. Before use in this study, *U2af1*(S34F) KI mice of mixed background (1) were serially backcrossed to C57BL/6 mice to achieve 100% C57BL/6 background using the Jackson Laboratory custom panel of 150 polymorphic SNPs. C57BL/6-CD45.1 mice were bred to C57BL/6-CD45.2 mice to generate C57BL/6-CD45.1/45.2 mice as a source of competitor bone marrow cells for competitive repopulation transplantation assays. Donor bone marrow from 7-12 weeks-old male and female mice were used for the experiments, and 8-week-old male and female C57BL/6-CD45.1 (Charles River) mice were used as recipients for the bone marrow transplantation assay. No gender bias was noted. Deletion of the *U2af1* KO allele and expression of *U2af1*(S34F) KI was achieved by six doses (300µg/mouse/dose) of polyinosinic-polycytidylic acid (Sigma) given every other day by intraperitoneal injection, or by six doses of tamoxifen (Sigma) dissolved in corn oil given by oral gavage, Monday, Wednesday, Friday, for two consecutive weeks (4mg/mouse/dose). The expression of transgenic U2AF1(WT) or U2AF1(S34F) was induced by Modified LabDiet® 5053 rodent chow with doxycycline (TestDiet).

**Apoptosis assay**

Apoptosis was assessed by flow cytometry using Annexin V. After staining with other antibodies as described, cells were washed and incubated in binding buffer containing Annexin V-APC and

SYTOX™ Green (Thermo Fisher) followed by flow cytometry analysis within an hour of staining.

**Western Blot**

Bulk bone marrow was lysed and sonicated in RIPA buffer (Cell Signaling) and protease inhibitor cocktail (Roche). Then, 10 μg of the protein sample was resolved in 10% SDS-PAGE gel (Bio-Rad) and transferred to a nitrocellulose membrane (Millipore). Protein was probed by anti-U2AF1 (Abcam) and anti-β-Actin (Sigma) antibodies. Detection was done with horseradish-peroxidase conjugated secondary mouse or rabbit antibodies (Cell Signaling), using SuperSignal West Pico and Femto chemiluminescence substrate (Thermo Scientific).

**RNA-Sequencing analysis**

Sequence read quality of the Illumina NovaSeq 6000-sequenced samples were assessed using FastQC [Andrews S. (2010). FastQC: a quality control tool for high throughput sequence data. Available online at http://www.bioinformatics.babraham.ac.uk/projects/fastqc/]. Consensus sequence logos or skipped exons were created using seqLogo (version 1.48.0 [Bembom O (2019). *seqLogo: Sequence logos for DNA sequence alignments*. R package version 1.48.0.]). Volcano plots were generated using enhanced volcano plot R package (version 1.6.0 [Blighe K, Rana S, Lewis M (2020). *EnhancedVolcano: Publication-ready volcano plots with enhanced colouring and labeling*. R package version 1.6.0.]) (2). We used the enrichment map function from clusterProfiler to organize the top 100 statistically significant mutually overlapping gene sets into clusters that share a common function.

### Supplemental figure legends

**Figure S1. Generation of a conditional *U2af1* knock-out allele.** (A) Targeting strategy to insert loxP sites flanking *U2af1* exon 2. (B-C) Successful targeting of five embryonic stem cell clones was verified by PCR and Southern blotting after digestion with the restriction enzyme MfeI. Wild-type C57BL/6 (B6) DNA was used as a control, and clone 243 was used to generate the *U2af1* knock-out mouse. (D) Whole-genome sequencing was performed on a heterozygous knock-out (*U2af1*$^{wt/flox}$) mouse. The heterozygous *U2af1*$^{wt/flox}$ mouse has sequence coverage for both *U2af1* wild-type allele and the targeted allele containing the LoxP sequences flanking exon 2, validating correct targeting of *U2af1* in the *U2af1*$^{wt/flox}$ mouse. LA (long arm), MA (middle arm), SA (short arm).

**Figure S2. Embryonic deletion of *U2af1* in hematopoietic cells is lethal and reduces the number of myeloid, lymphoid, and hematopoietic stem and progenitor cells.** (A) *U2af1*$^{flox/flox}$ mice were intercrossed with *Vav1*-Cre/*U2af1*$^{wt/flox}$ mice, and frequency of observed versus expected genotypes were analyzed at birth (Chi squared=13.158, 3 degrees of freedom, two-tailed p=0.0043, n=38). (B) Comparison of fetal liver development of embryos at embryonic day 14.5 (E14.5). (C) Absolute numbers of hematopoietic myeloid progenitor cells (n=3-10). (D) Apoptotic cells as a percent of Annexin V$^{+}$/ SYTOX Green$^{-}$ myeloid progenitor cells (n=3-10). All data are presented as mean +/- SD. *p<0.05, ***p<0.001, by one-way ANOVA with Tukey's multiple comparison test.

**Figure S3. *U2af1* deletion induces multi-lineage bone marrow failure.** (A) Experimental design of a non-competitive transplant of whole bone marrow cells from CD45.2 *Mx1*-Cre,

*U2af1*$^{flox/flox}$, *Mx1*-Cre/*U2af1*$^{wt/flox}$, or *Mx1*-Cre/*U2af1*$^{flox/flox}$ mice transplanted into lethally irradiated congenic wild-type CD45.1 recipient mice, followed by pIpC-induced Cre-activation and *U2af1* deletion. Analysis of the peripheral blood and bone marrow was done 8-11 days post-pIpC. (B) Survival curve analyzed by Mantel-Cox log-rank test, n=5 per group. (C) Spleen cellularity of *Mx1*-Cre, *U2af1*$^{wt/flox}$, and *Mx1*-Cre/*U2af1*$^{wt/flox}$ mice compared to *Mx1*-Cre/*U2af1*$^{flox/flox}$ mice 8-11 days post pIpC. (D) Total number of neutrophils, monocytes, B cells, and T cells per spleen at 8-11 days post pIpC for *Mx1*-Cre, *U2af1*$^{wt/flox}$, and *Mx1*-Cre/*U2af1*$^{wt/flox}$ mice compared to *Mx1*-Cre/*U2af1*$^{flox/flox}$ mice. (E). Number of progenitor (KL and KLS) cells per spleen of pIpC-treated mice transplanted with *Mx1*-Cre, *U2af1*$^{wt/flox}$, and *Mx1*-Cre/*U2af1*$^{wt/flox}$ bone marrow cells compared to *Mx1*-Cre/*U2af1*$^{flox/flox}$ bone marrow cells 8-11 days post pIpC. All data are presented as mean +/- SD. *p<0.05, **p<0.01, ***p<0.001, by one-way ANOVA with Tukey's multiple comparison test, n=5 per genotype.

**Figure S4. Hematopoietic stem cells are dependent on *U2af1* expression for survival.** (A) Experimental design of a competitive transplant of whole bone marrow cells from CD45.2 *Mx1*-Cre, *U2af1*$^{flox/flox}$, *Mx1*-Cre/*U2af1*$^{wt/flox}$, or *Mx1*-Cre/*U2af1*$^{flox/flox}$ mice mixed at a 1:1 ratio with congenic wild-type CD45.1/45.2 competitor cells followed by transplantation into lethally irradiated congenic wild-type CD45.1 recipient mice followed by pIpC-induced *U2af1* deletion. Peripheral blood and bone marrow chimerism of *Mx1*-Cre, *U2af1*$^{flox/flox}$, and *Mx1*-Cre/*U2af1*$^{wt/flox}$ mice was compared to *Mx1*-Cre/*U2af1*$^{flox/flox}$ mice. (B) Spleen chimerism of mature hematopoietic cells (neutrophils, monocytes, B cells, and T cells) at ten months post-pIpC (n=8). (C) Bone marrow chimerism of myeloid progenitor cells ten months post-pIpC (n=8 for KL and KLS; n=3-4 for all other). All data are presented as mean +/- SD. *p<0.05, ***p<0.001, by one-way ANOVA with Tukey's (B) or Dunnett's (C) multiple comparison test.

**Figure S5. *U2af1* deletion dysregulates genes involved in splicing, unfolded protein response, and apoptosis pathways.** Differential gene expression from RNA-seq of E14.5 hematopoietic progenitor cells (Lineage$^-$, cKit$^+$, Sca1$^-$). (A) *U2af1* exon 2 expression levels, counts per million (CPM) was calculated using deepTools (3). (B) Unsupervised hierarchical clustering of expressed genes. (C-D) Gene ontology (GO) enrichment analysis of the differentially regulated biological processes in *Vav1*-Cre/*U2af1*$^{\text{flox/flox}}$ compared to *U2af1*$^{\text{flox/flox}}$ control (group 1) and *Vav1*-Cre/*U2af1*$^{\text{wt/flox}}$ (group 2), and *U2af1*$^{\text{flox/flox}}$ control compared to *Vav1*-Cre/*U2af1*$^{\text{wt/flox}}$ (group 3); biological processes significantly upregulated (C) or downregulated (D) in *Vav1*-Cre/*U2af1*$^{\text{flox/flox}}$ cells, n=3 per genotype. *Gene pathways displayed in Figure 5.

**Figure S6. Survival of mutant U2AF1(S34F) hematopoietic cells is dependent on the expression of the residual wild-type allele and the ratio of U2AF1(WT:S34F) expression.** (A) Experimental design of a competitive transplant of whole bone marrow cells from CD45.2 tgU2AF1(WT)/*U2af1*$^{\text{wt/wt}}$, tgU2AF1(WT)/*U2af1*$^{\text{flox/flox}}$, tgU2AF1(S34F)/*U2af1*$^{\text{wt/wt}}$, or tgU2AF1(S34F)/*U2af1*$^{\text{flox/flox}}$ mice. All the donor test mice have both *Mx1*-Cre and rtTA. Transgenic U2AF1(WT) and U2AF1(S34F) were induced by 625 ppm or 10,000 ppm doxycycline chow (dox chow) followed by pIpC-induced *U2af1* deletion after four weeks, and analysis of the peripheral blood chimerism was performed. (B, C) Peripheral blood B cell and T cells chimerism in tgU2AF1(WT)/*U2af1*$^{\text{wt/wt}}$, tgU2AF1(WT)/*U2af1*$^{\text{flox/flox}}$, tgU2AF1(S34F)/*U2af1*$^{wt/wt}$, and tgU2AF1(S34F)/*U2af1*$^{flox/flox}$ mice after induction of transgenic U2AF1(WT) or U2AF1(S34F) by 10,000 ppm dox chow and *U2af1* deletion induced by pIpC Cre-activation (n=7-8). (D) Peripheral blood chimerism in tgU2AF1(WT)/*U2af1*$^{wt/wt}$ and tgU2AF1(S34F)/*U2af1*$^{wt/wt}$ mice after induction of transgenic U2AF1(WT) or U2AF1(S34F) by 625 ppm or 10,000 ppm dox chow and *U2af1* deletion induced by pIpC Cre-activation (n=7-8). (E) Experimental design of a competitive transplant of whole bone marrow cells from CD45.2 *Mx1*-Cre/rtTA/tgU2AF1(WT)/*U2af1*$^{wt/S34F}$ (tgU2AF1[WT]/*U2af1*$^{wt/S34F}$) mice. Transgenic U2AF1(WT) was induced by 10,000 ppm dox chow two weeks post-transplant followed by pIpC-induced *U2af1* deletion after two weeks of dox chow and analysis of peripheral blood chimerism was performed. (F, G) Peripheral blood B cell and T cell chimerism of tgU2AF1(WT)/*U2af1*$^{wt/S34F}$ with or without dox chow treatment (n=9-10). All data are presented as mean +/- SD. **p<0.01, ***p<0.001, by two-way ANOVA with Sidak's (F, G) or Tukey's (B-D) multiple comparison test.

**Figure S7. Hematopoietic cancer cells expressing mutant U2AF1(S34F) are sensitive to decreased levels of the wild-type U2AF1 expression.** (A) Experimental design of a competitive transplant of test whole bone marrow cells from CD45.2 tgU2AF1(WT)/*U2af1*$^{flox/flox}$ or tgU2AF1(WT)/*U2af1*$^{flox/S34F}$ mice mixed at a 1:1 ratio with congenic wild-type CD45.1/45.2 competitor cells followed by transplantation into lethally irradiated congenic wild-type CD45.1 recipient mice. All the donor test mice have both *Mx1*-Cre and rtTA. Transgenic U2AF1(WT) expression was induced by 10,000 ppm doxycycline chow (dox chow) two weeks post-transplant followed by pIpC induction of *U2af1* deletion two weeks later. Two weeks post-pIpC, the dox dose was reduced every two weeks (2500 ppm, 1250 ppm, 625 ppm, 350 ppm, 0 ppm), and analysis of the peripheral blood chimerism was performed. (B) A dose-finding curve showing peripheral blood neutrophil chimerism of tgU2AF1(WT)/*U2af1*$^{flox/flox}$ and tgU2AF1(WT)/*U2af1*$^{flox/S34F}$ (n=8-9), by two-way ANOVA with Tukey's multiple comparison test. (C) Western blot showing U2AF1 protein expression levels in bulk bone marrow cells of tgU2AF1(WT)/*U2af1*$^{wt/wt}$ mice after one week on doxycycline chow (0 ppm, 1250 ppm, 2500 ppm, or 10,000 ppm). (D) Experimental design of transplantation of tgU2AF1(WT)/*U2af1*$^{flox/S34F}$ MLL-AF9 AML tumor cells ($GFP^+$/ $CD45.2^+$) isolated from the spleen of primary mice into sub-lethally irradiated CD45.2 secondary recipients. Secondary recipients were treated with or without 10,000 ppm doxycycline chow followed by pIpC induction and analysis of the peripheral blood analysis and tumor watch. (E) $CD45.2^+$ MLL-AF9 AML cells chimerism up to 21 days post-second pIpC dose (n=10). (F) White blood cell count (WBC) up to 21 days post-second pIpC dose (n=10). (G) Spleen weight at the time of death (n=10). All data are presented as mean +/- SD. *p<0.05, **p<0.01, ***p<0.001, by one-way ANOVA with Tukey's (E) or Dunnett's (F, G) multiple comparison test.

### Supplemental Tables

**Supplemental Table 1: Differential gene expression data.** Submitted as a separate Excel file.

**Supplemental Table 2: GO enrichment analysis of differentially expressed genes.** Submitted as a separate Excel file.

**Supplemental Table 3: Differentially spliced junctions.** Submitted as a separate Excel file.

**Supplemental Table 4: Overlapping enrichment analysis result from GO, KEGG, and REACTOME on differentially spliced junctions**

| source | Term name | Term Id | Adjusted p-value |
|---|---|---|---|
| GO:CC | spliceosomal complex | GO:0005681 | 0.0410108 |
| KEGG | Spliceosome | KEGG:03040 | 0.022238 |
| REAC | mRNA Splicing | REAC:R-MMU-72172 | 0.0019292 |
| REAC | mRNA Splicing - Major Pathway | REAC:R-MMU-72163 | 0.0089023 |
| REAC | Processing of Capped Intron-Containing Pre-mRNA | REAC:R-MMU-72203 | 0.0188394 |

**Supplemental Table 5: Antibodies and other resources**

| Reagent or Resource | Source | Catalog number |
|---|---|---|
| Antibodies (clone) | | |
| Anti-CD45.1 BUV737 (A20) | BD Biosciences | Cat# 564574 |
| Anti-CD45.2 BUV395 (104) | BD Biosciences | Cat# 564616 |
| Anti-cKit BV421 (2B8) | Biolegend | Cat# 105828 |
| Anti-Sca-1 PE (D7) | Biolegend | Cat# 108108 |
| Anti-CD34 FITC (RAM34) | eBioscience | Cat# 11-0341-85 |
| Anti-CD16/32 BV711 (93) | Biolegend | Cat# 101337 |
| Anti-FLT3 APC (A2F10) | eBioscience | Cat# 17-1351-82 |
| Anti-CD48 PE-Cy7(HM48-1) | eBioscience | Cat# 25-0481-80 |
| Anti-CD150 APC/Fire 750 (TC15-12F12.2) | Biolegend | Cat# 115939 |
| Anti-CD45.1 BV421 (A20) | Biolegend | Cat# 110732 |
| Anti-CD45.2 FITC (104) | eBioscience | Cat# 11-0454-85 |
| Anti-CD115 PE (AFS98) | eBioscience | Cat# 12-1152-83 |
| Anti-B220 BV605 (RA3-6B2) | Biolegend | Cat# 103244 |
| Anti-CD3e APC (145-2C11) | eBioscience | Cat# 17-0031-83 |
| Anti-GR-1 APC/Fire 750 | Biolegend | Cat# 108456 |
| Anti-TER-119 BV605 (TER119) | Biolegend | Cat# 116239 |

| **Reagent or Resource** | **Source** | **Catalog number** |
|---|---|---|
| Anti-GR-1 BV605 (RB6-8C5) | Biolegend | Cat# 108440 |
| Anti-CD3e BV605 (145-2C11) | Biolegend | Cat# 100351 |
| Anti-CD41 BV605 (MWReg30) | Biolegend | Cat# 133921 |
| Anti-CD45.1 FITC (A20) | eBioscience | Cat# 11-0453-82 |
| Anti-CD45.2 APC (104) | eBioscience | Cat# 17-0454-82 |
| Anti-U2AF35/U2AF1 | Abcam | Cat# ab86305 |
| Anti-U2AF35/U2AF1 | Abcam | Cat# ab86305 |
| Anti-β-Actin (AC-15) | Sigma | Cat# A5441 |
| Bacterial and Virus Strains | | |
| MSCV MLL-AF9-IRES-GFP | Krivtsov et al. 2006 (4) | N/A |
| Chemicals, Peptides, and Recombinant Proteins | | |
| Polybrene | American Bioanalytical | Cat# AB01643-00001 |
| Polyinosinic-polycytidylic acid (pIpC) | Sigma | Cat# P1530 |
| Tamoxifen | Sigma | Cat# T5648 |
| Recombinant murine IL-3 | PeproTech | Cat# 213-13 |
| Recombinant murine SCF | PeproTech | Cat# 250-03 |
| Recombinant murine TPO | PeproTech | Cat# 315-14 |
| Recombinant murine FLT3 | PeproTech | Cat# 250-31L |
| Doxycycline chow | Test Diet | N/A |
| Critical Commercial Assays | | |
| MethoCult GF M3434 | StemCell Technologies | Cat# M3434 |
| KAPA RNA Hyper Prep Kit with RiboErase | KapaBiosystems | Cat# KR1351 |
| Experimental Models: Organisms/Strains | | |
| Mice: *Mx1*-Cre: B6.Cg-Tg(Mx1-cre)1Cgn/J | Jackson Laboratory | Cat# 003556 |
| Mice: *Vav1*-Cre: B6.Cg-$Commd10^{Tg(Vav1-icre)A2Kio}$/J | Jackson Laboratory | Cat# 008610 |
| Mice: Cre-*ERT2*: B6.Cg-Tg(CAG-cre/Esr1*)5Amc/J | Jackson Laboratory | Cat# 004682 |
| Mice: Transgenic U2AF1(S34F): B6.Cg-$Gt(ROSA)26Sor^{tm1(rtTA*M2)Jae}$ $Col1a1^{tm1(tetO-U2AF1*S34F)Mjwa}$/J | Shirai et al. 2015 (5), Jackson Laboratory | Cat# 029784 |
| Mice: Transgenic U2AF1(WT): B6.Cg-$Gt(ROSA)26Sor^{tm1(rtTA*M2)Jae}$ $Col1a1^{tm1(tetO-U2AF1*WT)Mjwa}$ | Shirai et al. 2015 (5) | N/A |
| Mice: *U2af1*(S34F) KI: $U2af1^{wt/S34F}$ | Fei et al. 2018 (1) | N/A |
| Mice: *U2af1* KO: $U2af1^{flox/flox}$ | This paper | N/A |
| Oligonucleotides | | |
| *U2af1* KO genotyping primer, forward: GAGGGTGATGGACCTACCCA | This paper | N/A |
| *U2af1* KO genotyping primer, reverse: GAGAAGAAAGGGGCGGCAAA | This paper | N/A |

| Reagent or Resource | Source | Catalog number |
|---|---|---|
| *U2af1* KO P1 primer: TCGTTCGAACATAACTTCGTATAG | This paper | N/A |
| *U2af1* KO P2 primer: TCTCCAACTGTGGGACTTACAGTG | This paper | N/A |
| *U2af1* KO Southern blot probe: ATCCTGTGGGAGCTAGAGTGGTGACCTCTAACACTGCCTGCAAGAGCTGCCTGACTTCTGTAGCATGGAAGAAAGTAGCTGAGAAGTAGCTTCAAGTTAATGGCGTTTCTGCCACATCTTGCCAGAAATAATCAGTTCTTGTTTTCCCTTTACAGAGTCAACTGTTCATTTTATTTCAAAATCGGAGCATGTCGTCATGGAGACAGATGTTCTCGGTTGCACAATAAACCAACCTTTAGCCAGGTTTGTGTGTTGCTGGGTTTTTTGCTTTTTTTTTTTTTATTTTTCACAAATTATCAAAAGTTCTTGTGCTTTCAGGAGCGATTAACATTCTCATGCACTCAGCAAGTTAAATCTTCCCTCTGGTTTATTTTGCATTTGTTACAGAGGGTGATGGACCTACCCAGTGGGAAGTCTTGA | This paper | N/A |

### Figure 1S

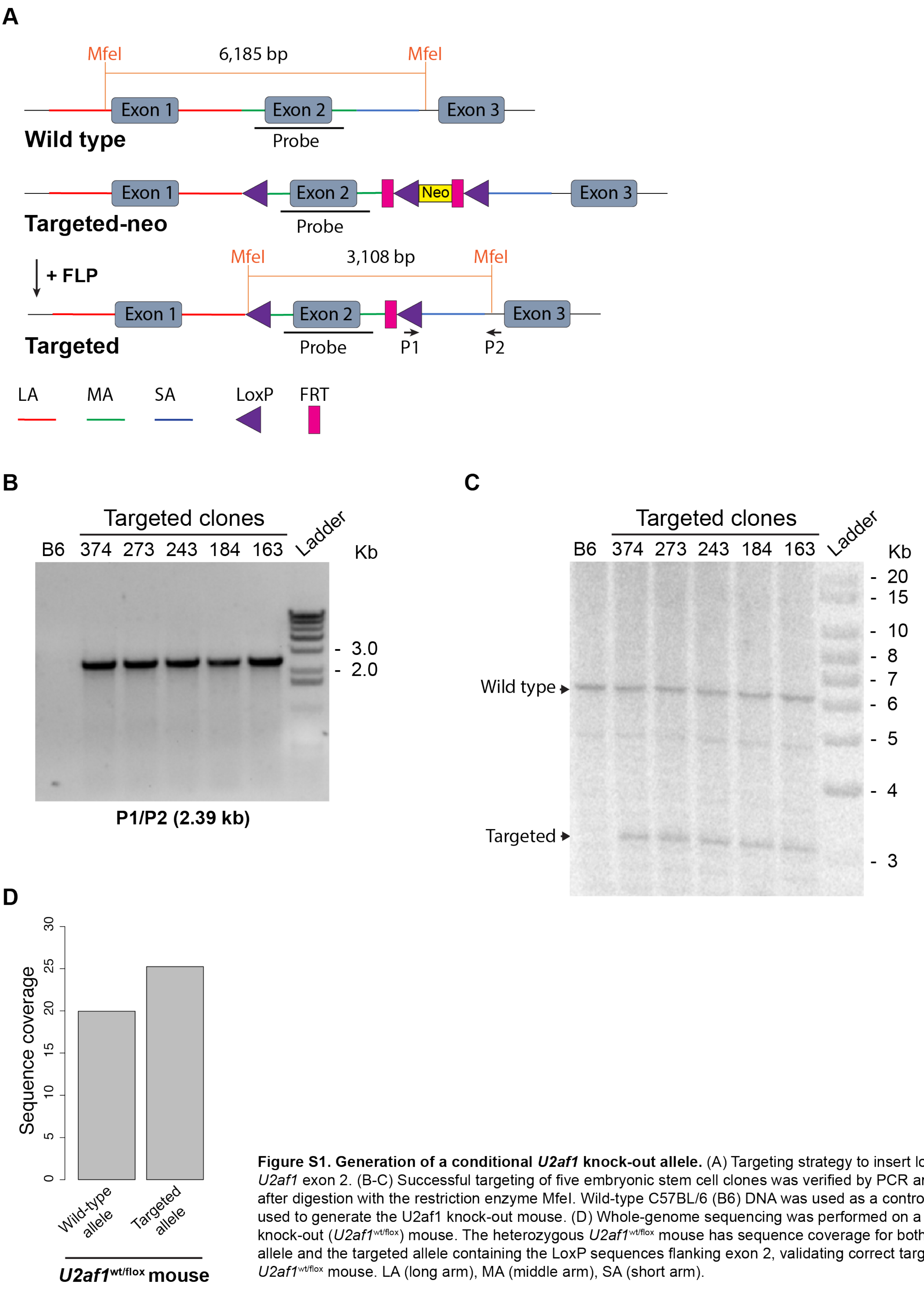


**Figure S1. Generation of a conditional *U2af1* knock-out allele.** (A) Targeting strategy to insert loxP sites flanking *U2af1* exon 2. (B-C) Successful targeting of five embryonic stem cell clones was verified by PCR and Southern blotting after digestion with the restriction enzyme MfeI. Wild-type C57BL/6 (B6) DNA was used as a control, and clone 243 was used to generate the U2af1 knock-out mouse. (D) Whole-genome sequencing was performed on a heterozygous knock-out (*U2af1*$^{wt/flox}$) mouse. The heterozygous *U2af1*$^{wt/flox}$ mouse has sequence coverage for both *U2af1* wild-type allele and the targeted allele containing the LoxP sequences flanking exon 2, validating correct targeting of *U2af1* in the *U2af1*$^{wt/flox}$ mouse. LA (long arm), MA (middle arm), SA (short arm).

### Figure S2

**A**

| | | | | |
|---|---|---|---|---|
| ***Vav1*-Cre** | - | - | + | + |
| ***U2af1*** | wt/flox | flox/flox | wt/flox | flox/flox |
| **Live Births (n=38)** | 13 (34.2%) | 11 (28.9%) | 14 (36.8%) | 0 |
| **Expected** | 25% | 25% | 25% | 25% |

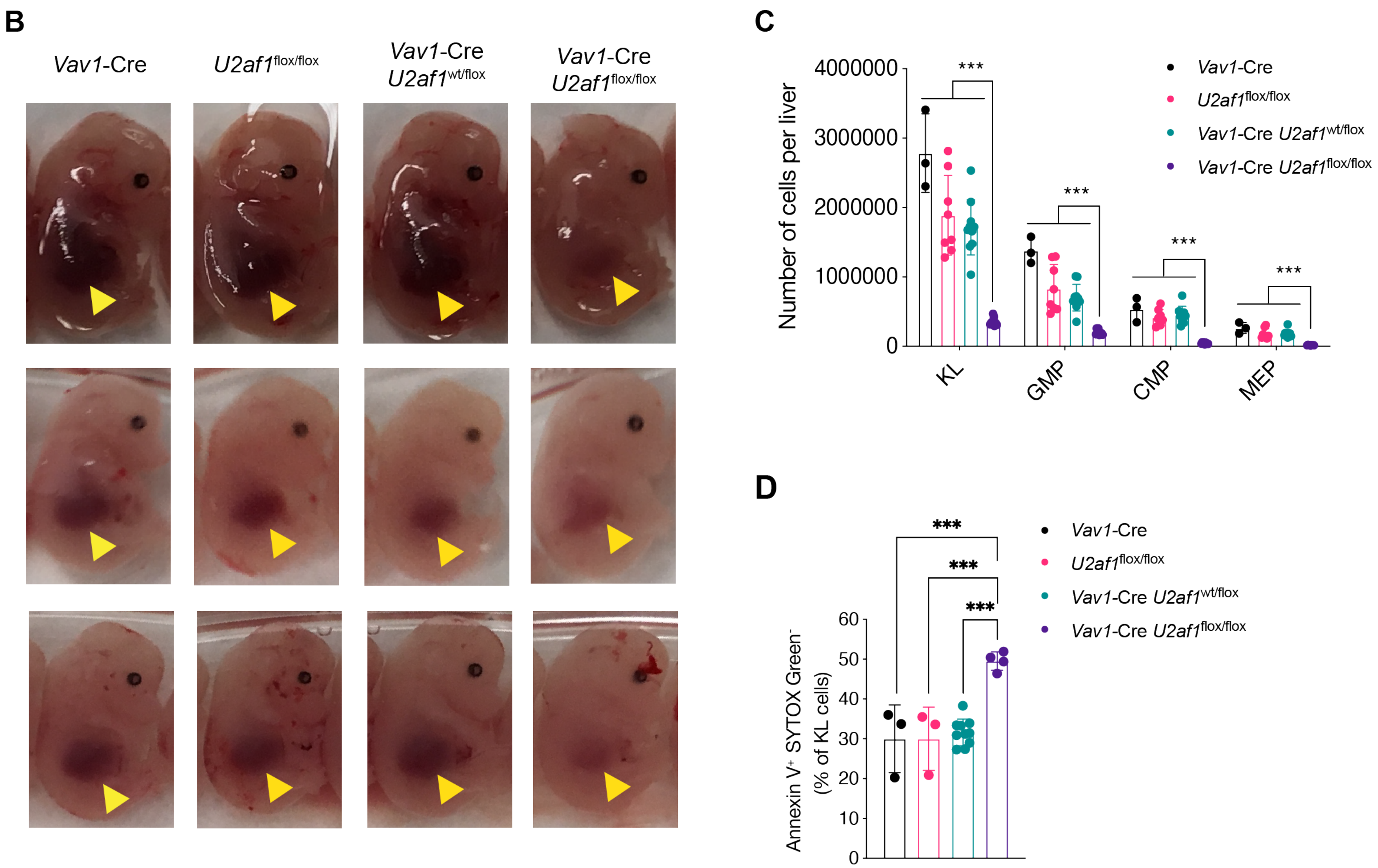


**Figure S2. Embryonic deletion of *U2af1* in hematopoietic cells is lethal and reduces the number of myeloid, lymphoid, and hematopoietic stem and progenitor cells.** (A) *U2af1*$^{flox/flox}$ mice were intercrossed with *Vav1*-Cre/*U2af1*$^{wt/flox}$ mice, and frequency of observed versus expected genotypes were analyzed at birth (Chi squared=13.158, 3 degrees of freedom, two-tailed p=0.0043, n=38). (B) Comparison of fetal liver development of embryos at embryonic day 14.5 (E14.5). (C) Absolute numbers of hematopoietic myeloid progenitor cells (n=3-10). (D) Apoptotic cells as a percent of Annexin V$^+$/ SYTOX Green$^-$ myeloid progenitor cells (n=3-10). All data are presented as mean +/- SD. *p<0.05, ***p<0.001, by one-way ANOVA with Tukey's multiple comparison test.

### Figure S3

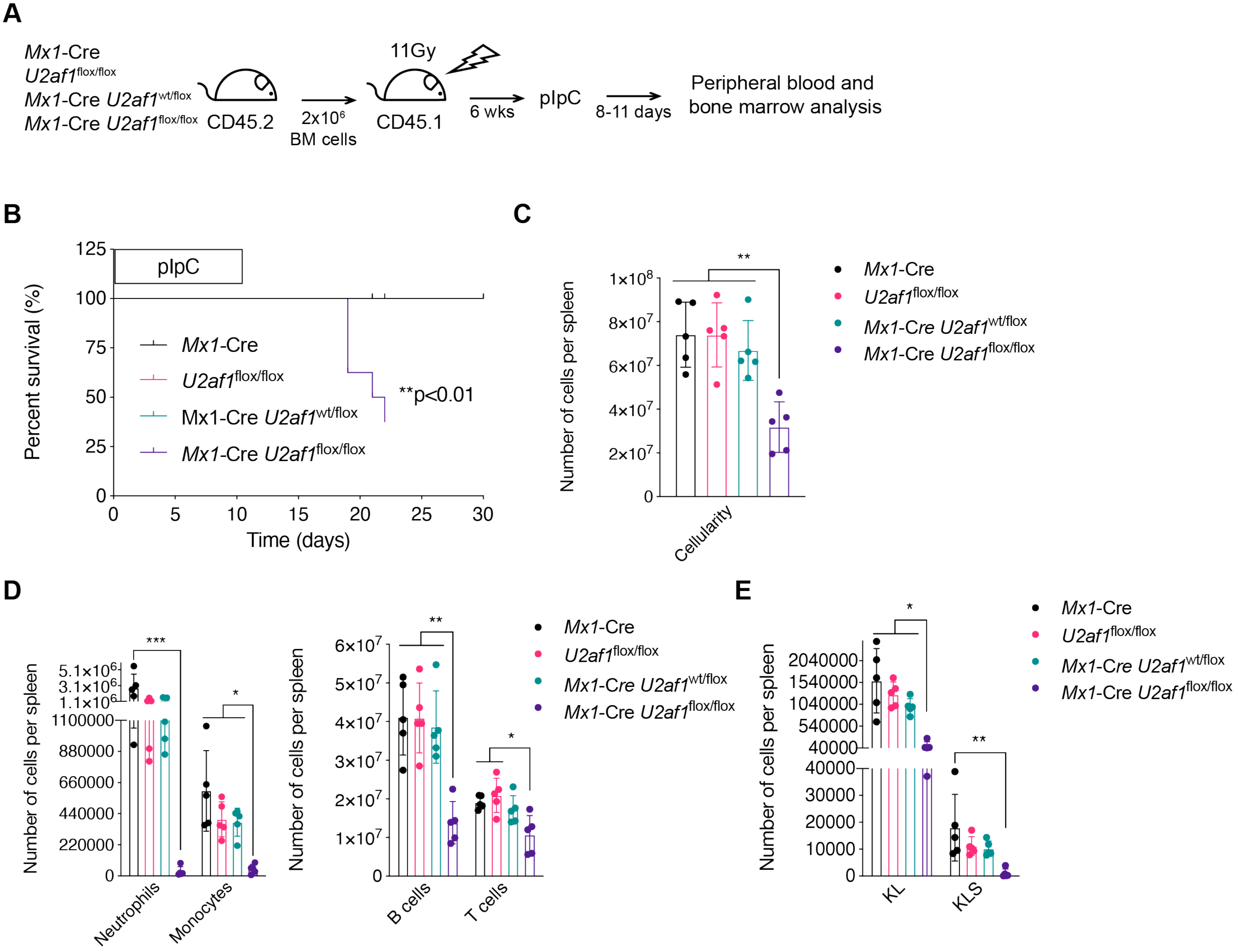


**Figure S3. *U2af1* deletion induces multi-lineage bone marrow failure.** (A) Experimental design of a non-competitive transplant of whole bone marrow cells from CD45.2 Mx1-Cre, *U2af1*$^{flox/flox}$, *Mx1*-Cre/*U2af1*$^{wt/flox}$, or *Mx1*-Cre/*U2af1*$^{flox/flox}$ mice transplanted into lethally irradiated congenic wild-type CD45.1 recipient mice, followed by pIpC-induced Cre-activation and *U2af1* deletion. Analysis of the peripheral blood and bone marrow was done 8-11 days post-pIpC. (B) Survival curve analyzed by Mantel-Cox log-rank test, n=5 per group. (C) Spleen cellularity of *Mx1*-Cre, *U2af1*$^{wt/flox}$, and *Mx1*-Cre/*U2af1*$^{wt/flox}$ mice compared to *Mx1*-Cre/*U2af1*$^{flox/flox}$ mice 8-11 days post pIpC. (D) Total number of neutrophils, monocytes, B cells, and T cells per spleen at 8-11 days post pIpC for *Mx1*-Cre, *U2af1*$^{wt/flox}$, and *Mx1*-Cre/*U2af1*$^{wt/flox}$ mice compared to *Mx1*-Cre/*U2af1*$^{flox/flox}$ mice. (E). Number of progenitor (KL and KLS) cells per spleen of pIpC-treated mice transplanted with *Mx1*-Cre, *U2af1*$^{wt/flox}$, and *Mx1*-Cre/*U2af1*$^{wt/flox}$ bone marrow cells compared to *Mx1*-Cre/*U2af1*$^{flox/flox}$ bone marrow cells 8-11 days post pIpC. All data are presented as mean +/- SD. *p<0.05, **p<0.01, ***p<0.001, by one-way ANOVA with Tukey's multiple comparison test, n=5 per genotype.

### Figure S4

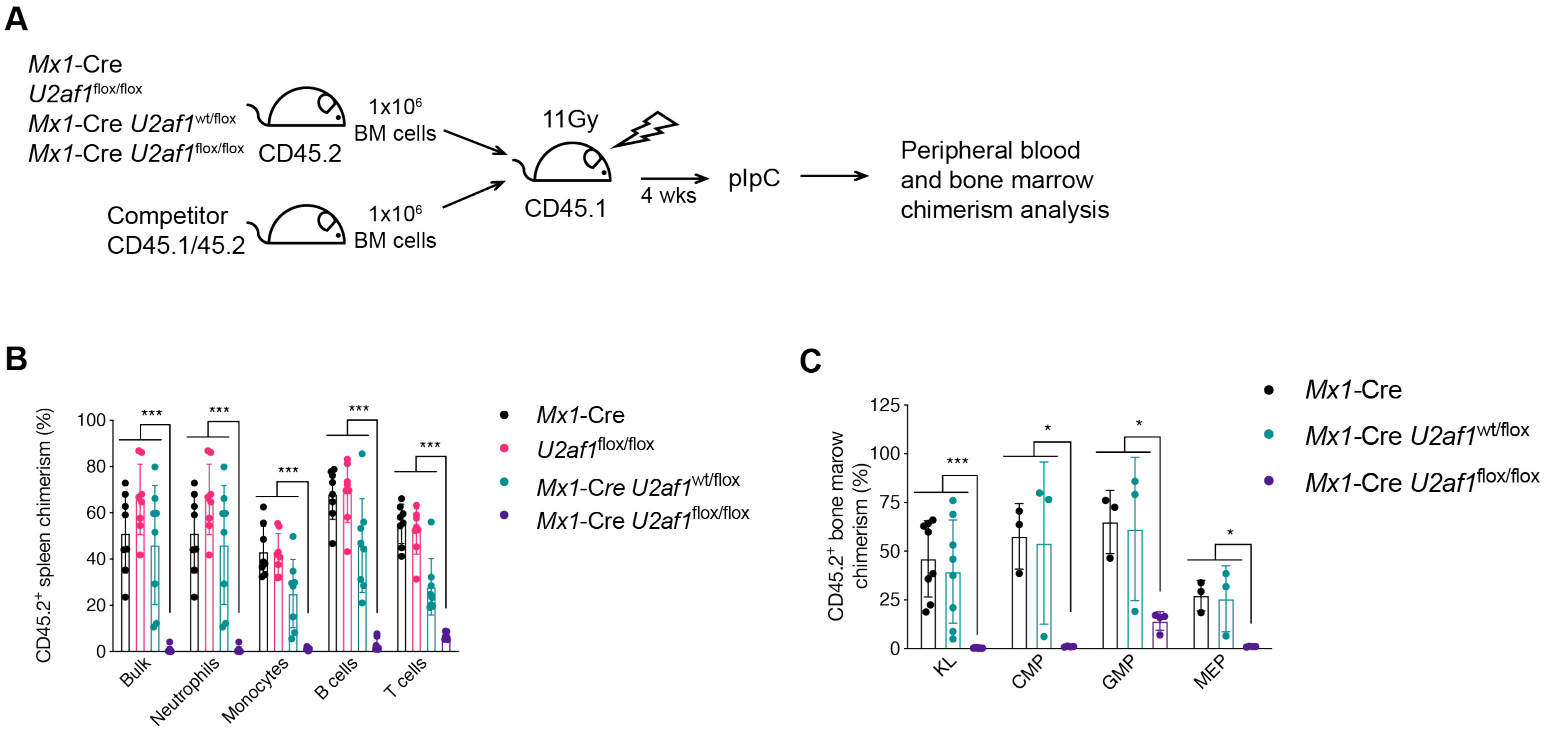


**Figure S4. Hematopoietic stem cells are dependent on *U2af1* expression for survival.** (A) Experimental design of a competitive transplant of whole bone marrow cells from CD45.2 *Mx1*-Cre, *U2af1*$^{flox/flox}$, *Mx1*-Cre/*U2af1*$^{wt/flox}$, or *Mx1*-Cre/*U2af1*$^{flox/flox}$ mice mixed at a 1:1 ratio with congenic wild-type CD45.1/45.2 competitor cells followed by transplantation into lethally irradiated congenic wild-type CD45.1 recipient mice followed by pIpC-induced *U2af1* deletion. Peripheral blood and bone marrow chimerism of *Mx1*-Cre, *U2af1*$^{flox/flox}$, and *Mx1*-Cre/*U2af1*$^{wt/flox}$ mice was compared to *Mx1*-Cre/*U2af1*$^{flox/flox}$ mice. (B) Spleen chimerism of mature hematopoietic cells (neutrophils, monocytes, B cells, and T cells) at ten months post-pIpC (n=8). (C) Bone marrow chimerism of myeloid progenitor cells ten months post-pIpC (n=8 for KL and KLS; n=3-4 for all other). All data are presented as mean +/- SD. *p<0.05, ***p<0.001, by one-way ANOVA with Tukey's (B) or Dunnett's (C) multiple comparison test.

**Figure S5**

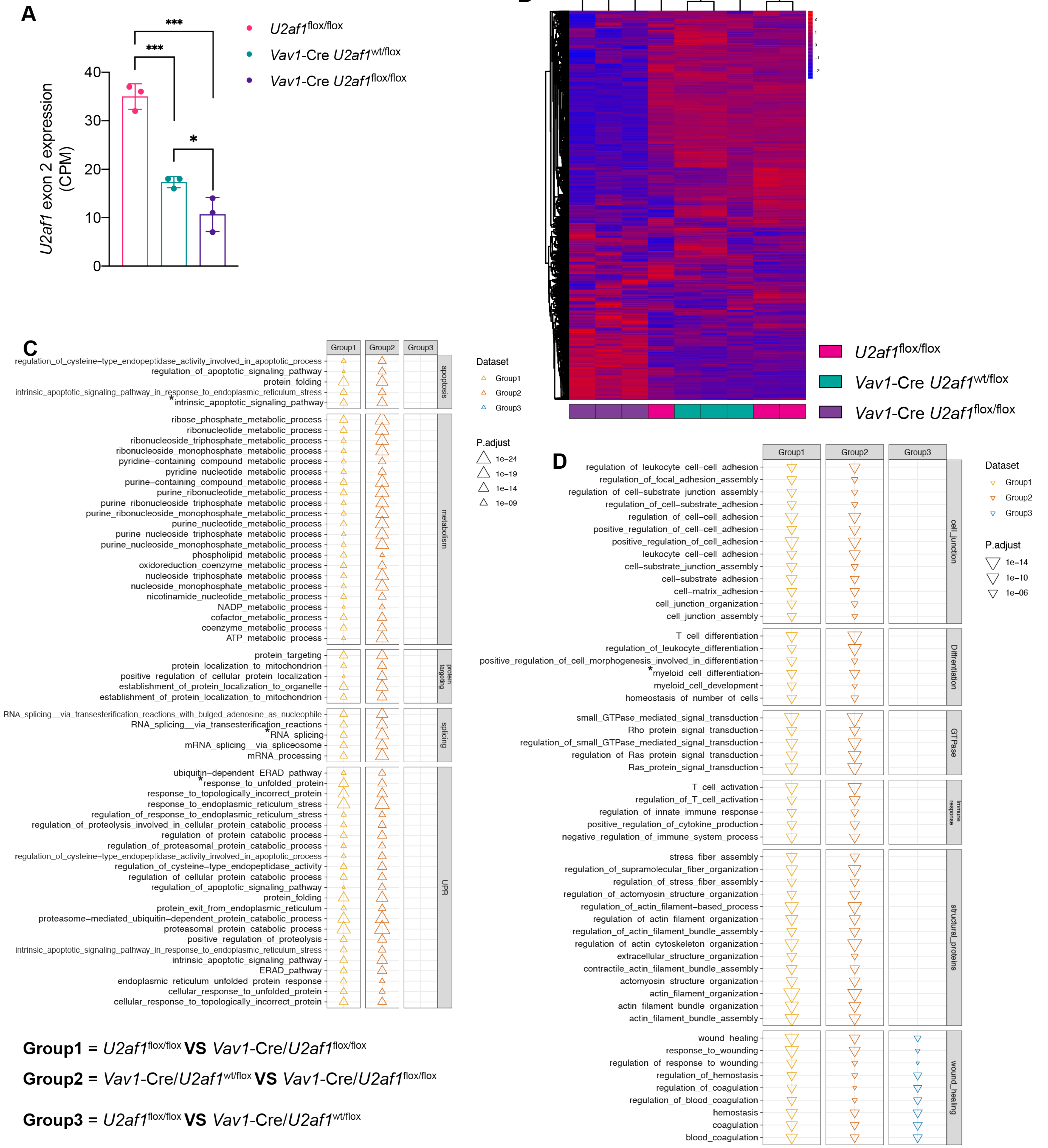


**Figure S5. *U2af1* deletion dysregulates genes involved in splicing, unfolded protein response, and apoptosis pathways.** Differential gene expression from RNA-seq of E14.5 hematopoietic progenitor cells (Lineage-, cKit+, Sca1-). (A) ***U2af1*** exon 2 expression levels, counts per million (CPM) was calculated using deepTools (3). (B) Unsupervised hierarchical clustering of expressed genes. (C-D) Gene ontology (GO) enrichment analysis of the differentially regulated biological processes in *Vav1*-Cre/*U2af1*$^{flox/flox}$ compared to *U2af1*$^{flox/flox}$ control (group 1) and *Vav1*-Cre/*U2af1*$^{wt/flox}$ (group 2), and *U2af1*$^{flox/flox}$ control compared to *Vav1*-Cre/*U2af1*$^{wt/flox}$ (group 3); biological processes significantly upregulated (C) or downregulated (D) in *Vav1*-Cre/*U2af1*$^{flox/flox}$ cells, n=3 per genotype. *Gene pathways displayed in Figure 5.

### Figure S6

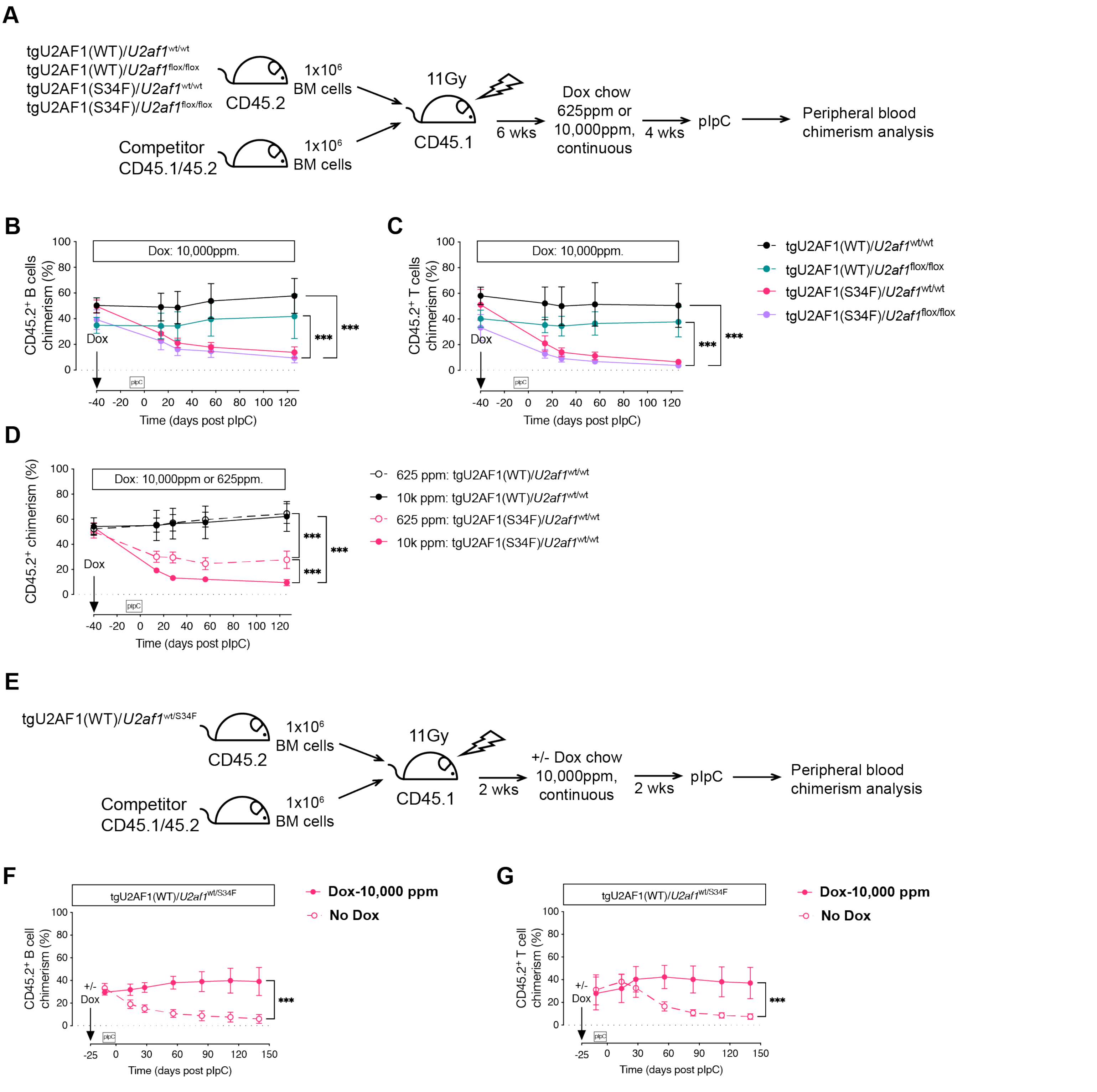


**Figure S6. Survival of mutant U2AF1(S34F) hematopoietic cells is dependent on the expression of the residual wild-type allele and the ratio of U2AF1(WT:S34F) expression.** (A) Experimental design of a competitive transplant of whole bone marrow cells from CD45.2 tgU2AF1(WT)/*U2af1*$^{wt/wt}$, tgU2AF1(WT)/*U2af1*$^{flox/flox}$, tgU2AF1(S34F)/*U2af1*$^{wt/wt}$, or tgU2AF1(S34F)/*U2af1*$^{flox/flox}$ mice. All the donor test mice have both Mx1-Cre and rtTA. Transgenic U2AF1(WT) and U2AF1(S34F) were induced by 625 ppm or 10,000 ppm doxycycline chow (dox chow) followed by pIpC-induced U2af1 deletion after four weeks, and analysis of the peripheral blood chimerism was performed. (B, C) Peripheral blood B cell and T cells chimerism in tgU2AF1(WT)/*U2af1*$^{wt/wt}$, tgU2AF1(WT)/*U2af1*$^{flox/flox}$, tgU2AF1(S34F)/*U2af1*$^{wt/wt}$, and tgU2AF1(S34F)/*U2af1*$^{flox/flox}$ mice after induction of transgenic U2AF1(WT) or U2AF1(S34F) by 10,000 ppm dox chow and *U2af1* deletion induced by pIpC Cre-activation (n=7-8). (D) Peripheral blood chimerism in tgU2AF1(WT)/*U2af1*$^{wt/wt}$ and tgU2AF1(S34F)/*U2af1*$^{wt/wt}$ mice after induction of transgenic U2AF1(WT) or U2AF1(S34F) by 625 ppm or 10,000 ppm dox chow and *U2af1* deletion induced by pIpC Cre-activation (n=7-8). (E) Experimental design of a competitive transplant of whole bone marrow cells from CD45.2 Mx1-Cre/rtTA/t-gU2AF1(WT)/*U2af1*$^{wt/S34F}$ (tgU2AF1[WT]/*U2af1*$^{wt/S34F}$) mice. Transgenic U2AF1(WT) was induced by 10,000 ppm dox chow two weeks post-transplant followed by pIpC-induced *U2af1* deletion after two weeks of dox chow and analysis of peripheral blood chimerism was performed. (F, G) Peripheral blood B cell and T cell chimerism of tgU2AF1(WT)/*U2af1*$^{wt/S34F}$ with or without dox chow treatment (n=9-10). All data are presented as mean +/- SD. **p<0.01, ***p<0.001, by two-way ANOVA with Sidak's (F, G) or Tukey's (B-D) multiple comparison test.

### Figure S7

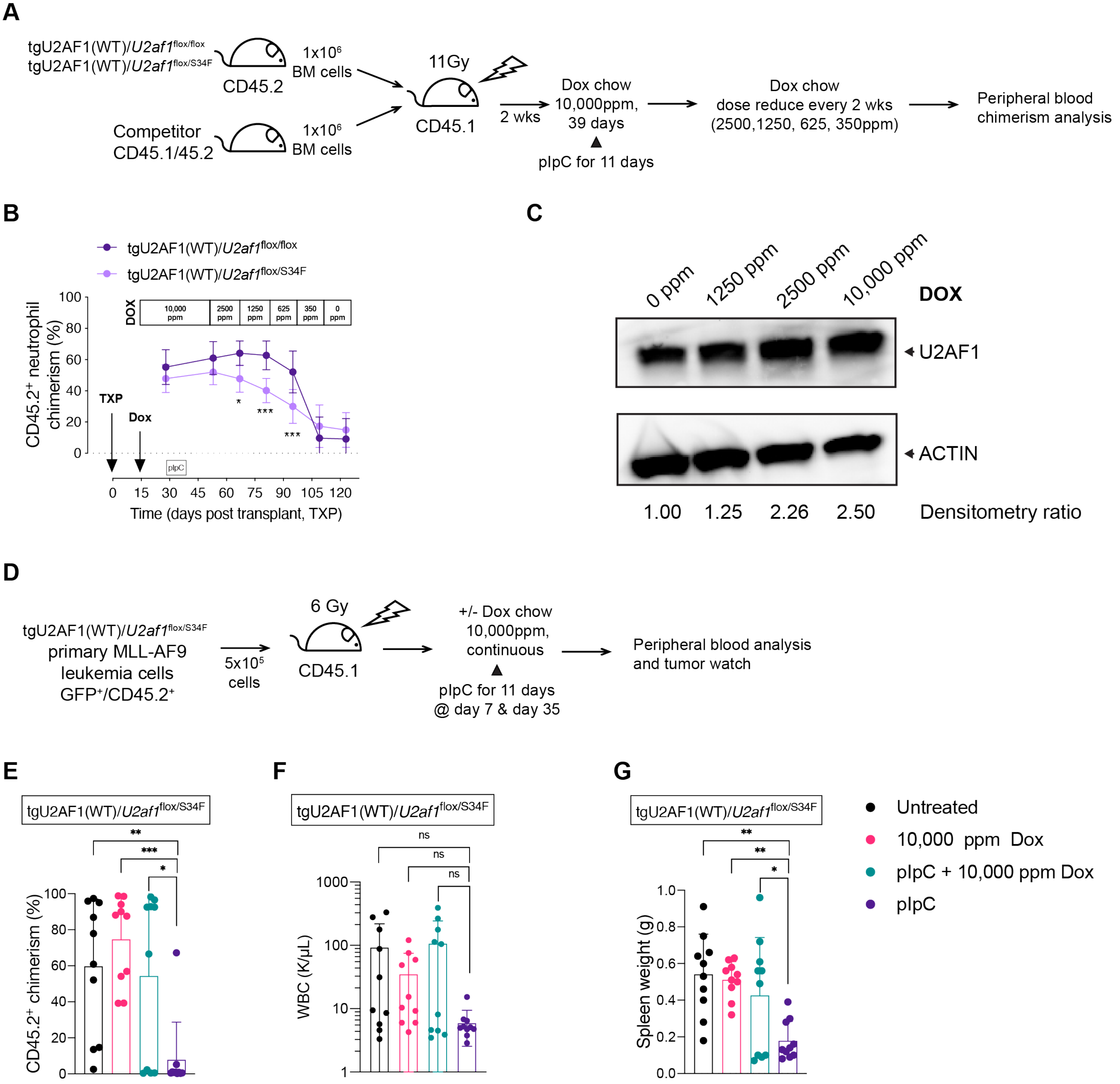


**Figure S7. Hematopoietic cancer cells expressing mutant U2AF1(S34F) are sensitive to decreased levels of the wild-type U2AF1 expression.** (A) Experimental design of a competitive transplant of test whole bone marrow cells from CD45.2 tgU2AF1(WT)/*U2af1*$^{flox/flox}$ or tgU2AF1(WT)/*U2af1*$^{flox/S34F}$ mice mixed at a 1:1 ratio with congenic wild-type CD45.1/45.2 competitor cells followed by transplantation into lethally irradiated congenic wild-type CD45.1 recipient mice. All the donor test mice have both *Mx1*-Cre and rtTA. Transgenic U2AF1(WT) expression was induced by 10,000 ppm doxycycline chow (dox chow) two weeks post-transplant followed by pIpC induction of *U2af1* deletion two weeks later. Two weeks post-pIpC, the dox dose was reduced every two weeks (2500 ppm, 1250 ppm, 625 ppm, 350 ppm, 0 ppm), and analysis of the peripheral blood chimerism was performed. (B) A dose-finding curve showing peripheral blood neutrophil chimerism of tgU2AF1(WT)/*U2af1*$^{flox/flox}$ and tgU2AF1(WT)/*U2af1*$^{flox/S34F}$ (n=8-9), by two-way ANOVA with Tukey's multiple comparison test. (C) Western blot showing U2AF1 protein expression levels in bulk bone marrow cells of tgU2AF1(WT)/*U2af1*$^{wt/wt}$ mice after one week on doxycycline chow (0 ppm, 1250 ppm, 2500 ppm, or 10,000 ppm). (D) Experimental design of transplantation of tgU2AF1(WT)/*U2af1*$^{flox/S34F}$ MLL-AF9 AML tumor cells (GFP+/ CD45.2+) isolated from the spleen of primary mice into sub-lethally irradiated CD45.2 secondary recipients. Secondary recipients were treated with or without 10,000 ppm doxycycline chow followed by pIpC induction and analysis of the peripheral blood analysis and tumor watch. (E) CD45.2+ MLL-AF9 AML cells chimerism up to 21 days post-second pIpC dose (n=10). (F) White blood cell count (WBC) up to 21 days post-second pIpC dose (n=10). (G) Spleen weight at the time of death (n=10). All data are presented as mean +/- SD. *p<0.05, **p<0.01, ***p<0.001, by one-way ANOVA with Tukey's (E) or Dunnett's (F, G) multiple comparison test.
